# The basal ganglia can control learned motor sequences independently of motor cortex

**DOI:** 10.1101/827261

**Authors:** Ashesh K. Dhawale, Steffen B. E. Wolff, Raymond Ko, Bence P. Ölveczky

## Abstract

How the basal ganglia contribute to the execution of learned motor skills has been thoroughly investigated. The two dominant models that have emerged posit roles for the basal ganglia in action selection and in the modulation of movement vigor. Here we test these models in rats trained to execute highly stereotyped and idiosyncratic task-specific motor sequences. Recordings and manipulations of neural activity in the striatum were not well explained by either model, and suggested that the basal ganglia, in particular its sensorimotor arm, are crucial for controlling the detailed kinematic structure of the learned behaviors. Importantly, the neural representations in the striatum, and the control functions they subserve, did not depend on the motor cortex. Taken together, these results extend our understanding of basal ganglia function, by suggesting that they can control and modulate lower-level subcortical motor circuits on a moment-by-moment basis to generate stereotyped learned motor sequences.

## Introduction

Much of what we do in our daily lives – be it tying our shoelaces or playing sports – relies on our brain’s ability to learn and execute stereotyped task-specific motor sequences^1^. The basal ganglia (BG), a collection of phylogenetically conserved midbrain structures^2–4^, have been implicated in their acquisition and proper execution^5–9^. Yet despite intense interest in deciphering BG function, their exact contributions to motor skill execution remains a matter of debate.

Two major models have emerged. One, which we refer to as the ‘vigor’ model^7, 10, 11^, proposes that the BG modulate the speed and amplitude (or ‘vigor’) of learned movements and sequential actions (Figure 1A). Observations that activity in the BG covaries with vigor^12–15^ and follows, rather than leads, movement initiation^16^, provides support for this model. Furthermore, manipulations of the BG in both primates^17, 18^ and rodents^12, 13, 15^ can affect movement vigor without overly influencing the sequential organization of the behaviors.

**Figure 1:**
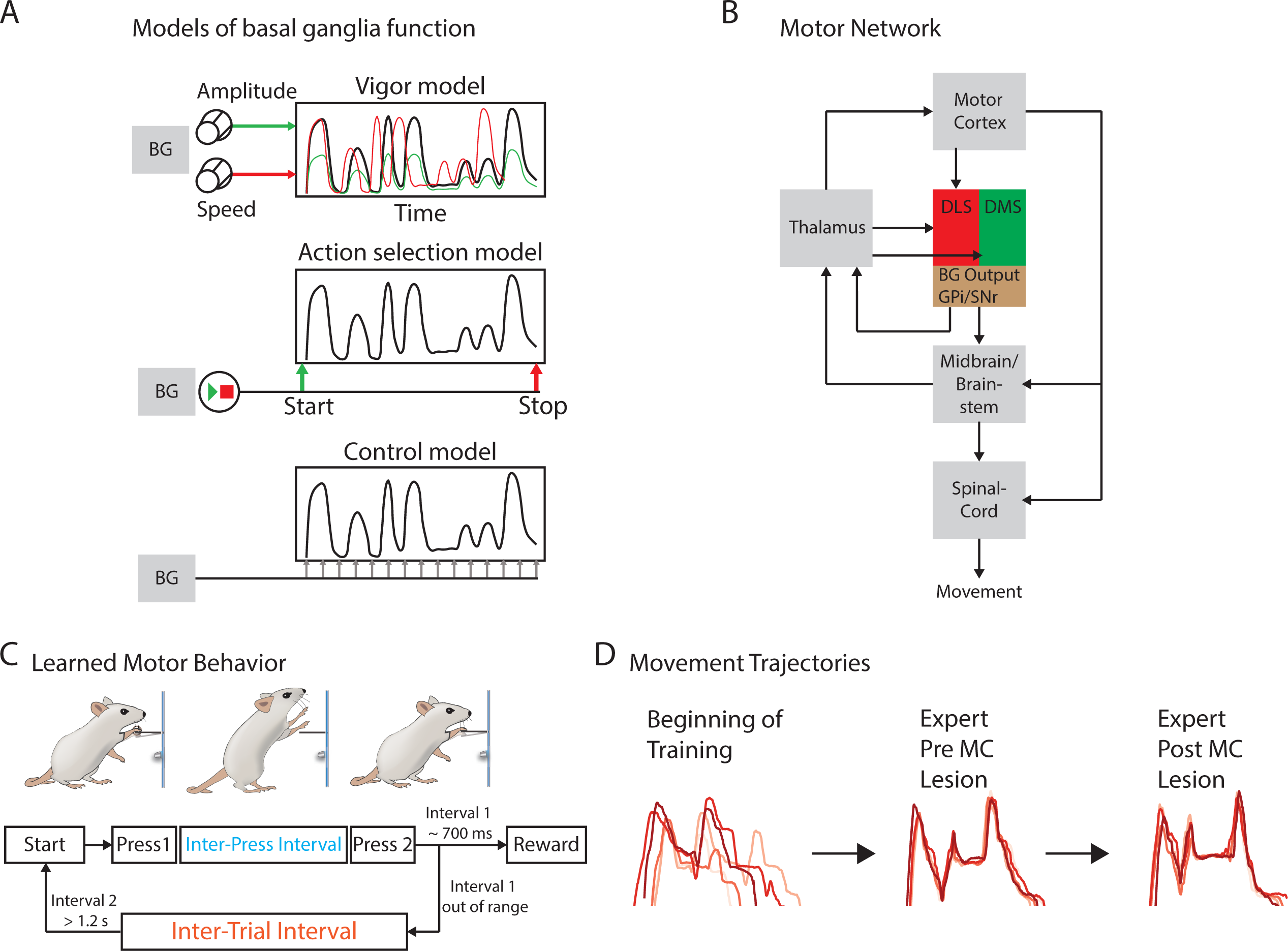
Probing the role of the basal ganglia (BG) in the execution of learned motor sequences. **A.** Simplified models for the function of the BG in the execution of learned motor skills. *Top*: The ‘vigor’ model suggests that the BG modulate parameters like the amplitude or speed of learned motor sequences, symbolized here by dials which can be turned by the BG. A learned motor sequence (black trace), which unfolds over time, can be changed in amplitude (green) or speed (red) or both (not shown), without changing the overall sequence or structure of the behavior. *Middle*: The ‘action selection’ model posits that the BG can select and then initiate (green) and terminate (red) learned motor sequences, appropriate for a given context, but does not affect its structure or kinematics. *Bottom*: An alternative way for the BG to contribute to the control of motor skills. Here the BG controls execution level detail on a moment-by moment basis. This can be seen as analogous to the assumed role of pre-motor nucleus HVC in songbirds (see text for details). **B.** Simple schematic of the motor circuits relevant to this study. The BG can affect the execution of learned behaviors via their influence on motor cortex, through the cortico-BG-thalamo-cortical loop, and/or via direct projections to the brainstem mid-brain motor centers. The dorsolateral striatum (DLS) receives much of its input from sensorimotor cortex, and defines the sensorimotor arm of the BG. **C.** Behavioral paradigm to probe the role of the BG in motor skill execution^31^. Rats are rewarded for pressing a lever twice with a specific target interval (inter-press interval – IPI). After unsuccessful trials, animals can only initiate a new trial after refraining from pressing the lever for a given interval (inter-trial interval – ITI). **D.** Over the course of training, animals develop stereotyped motor sequences that conform to the constraints of the task. These learned motor sequences are preserved in largely unaltered form after motor cortex lesions^31^.

The other main idea, which we call the ‘action selection’ model, posits that the main function of the BG is to select appropriate actions by providing start and stop commands to the downstream control circuits that enact them^8, 9, 19–21^ (Figure 1A). This view has received support from recordings in both rodents^22–25^ and primates^26^ showing that neural activity in striatum, GPe and SNr^27^ preferentially represents the initiation and termination of over-trained behaviors.

The common denominator of the two models is that the BG do not directly control the detailed structure of the learned behaviors^7^, but rather exert their influence on motor output by modulating or triggering the control circuits that do. In analogy to playing music on a jukebox, the BG have buttons for initiating and terminating a particular song (‘action selection’ model) and/or dials to control its volume and bass levels (‘vigor’ model), but no ability to affect its melody or lyrics.

For learned motor skills, the control circuit widely assumed to be the BG’s main target is the motor cortex (via thalamus, Figure 1B)^28–30^. However, a recent study showing that motor skills in rats can be executed without motor cortex^31^, suggests that lower-level circuits can be essential controllers for learned behaviors. Incidentally, the BG send direct projections to brainstem and midbrain motor centers^32–35^, including superior colliculus, periaqueductal gray, and various pontine and medullary reticular nuclei. These projections are part of the phylogenetically older ‘BG-subcortical pathway’, which is thought to be involved in selecting^32, 34–38^, sequencing^39–44^, and modulating^45, 46^ innate behaviors. Whether this BG-subcortical pathway, often thought of as a hardwired circuit for species-typical behaviors^34, 36, 37^, can assume a leading role in the execution of learned motor skills is not known.

A hint that this lower-level BG pathway may be involved comes from considering the motor cortex-independent skills alluded to above^31^ (Figure 1C,D). These behaviors are shaped by trial-and-error learning into highly idiosyncratic and task-specific motor sequences with rich and reproducible kinematic structure (Figure 1D). This would seem to require a degree of experience-dependent plasticity not typically associated with the brainstem and midbrain circuits. Striatum, the major input nucleus of the BG, however, receives dense dopaminergic innervation^47^, is a known player in reinforcement learning^48, 49^, and has the ability to influence control circuits through the BG’s output projections^33^. One possibility, then, is that the BG assume a control function and learn to orchestrate pattern generators in downstream motor circuits to produce new and adaptive motor sequences (‘control model’ in Figure 1A). This, however, would require us to extend the established models and theories relating how and what the BG contribute to motor skill execution. Thus, rather than thinking about BG as playing on an old jukebox, the more apt analogy would be that they function as a modern-day DJ, who can mix up new material to fit a particular situation.

To test this possibility, and probe the role of the BG more broadly, we utilized the behavioral paradigm mentioned above (Figure 1C) that results in motor skills robust to motor cortex lesions^31^ (Figure 1D). We focus our investigation primarily on the striatum, distinguishing its sensorimotor region (dorsolateral striatum, DLS), which receives input from sensorimotor cortex as well as thalamus ^50, 51^, and its associative region (dorsomedial striatum, DMS), which, in addition to thalamic input^50, 51^ receives projections from prefrontal and parietal cortices^50, 52, 53^.

Combining chronic neural recordings and high-resolution behavioral tracking, we find that activity in the DLS, but not in the DMS, represents execution-level details of the learned behaviors such as their temporal progression and kinematics, and does so even after removal of motor cortical input. Lesions of the DLS, but not the DMS, disrupted the task-specific motor sequences, reverting animals to behaviors expressed early in training. These results are not readily explained by existing models of the BG (as a ‘jukebox’, top two models in Figure 1A) and suggest that their function can extend beyond action selection and modulation of vigor to involve the moment-to-moment control of learned behavior (more like a ‘modern day DJ’, bottom model in Figure 1A). This function is likely instantiated through BG’s projections to brainstem and midbrain motor centers^32–35^ (Figure 1B), and is independent of, and cannot be subsumed by, motor cortex. Overall, these results extend our understanding of how the BG contribute to motor skill execution.

## Results

### The DLS is significantly more modulated than DMS during execution of a learned motor sequence

To probe whether and how the BG contribute to the execution of stereotyped learned motor sequences, we trained rats in our timed lever pressing task, in which a water reward is delivered contingent on animals pressing a lever twice separated by a specific time interval (inter-press interval or IPI ; target: 700 ms, see Methods) (Figure 1C)^31^. Over about a month of daily training, rats developed idiosyncratic and highly precise movement patterns (Figure 1D). Once acquired, these skilled behaviors are stably executed over long periods of time and robust to motor cortex lesions^31^ (Figure 1D).

We first sought to describe how neurons in the striatum represent these learned motor sequences. For this, we implanted expert rats with tetrode drives^54^. We had identified the DLS and the DMS by anterograde viral tracing from motor and prefrontal cortices respectively (Supplementary Figure 1) and targeted our recordings to these subregions in separate cohorts of animals (n=3 each for the DLS and the DMS, Figure 2A). We recorded from large populations of striatal neurons (in total, n=1591 units in DLS and n=1176 units in the DMS) continuously over several weeks of training^54^. Simultaneous with our neural recordings, we also monitored the animals’ movements using high-speed videography and automated markerless tracking of body parts such as the paws and head^55, 56^ (Figure 2A).

**Figure 2:**
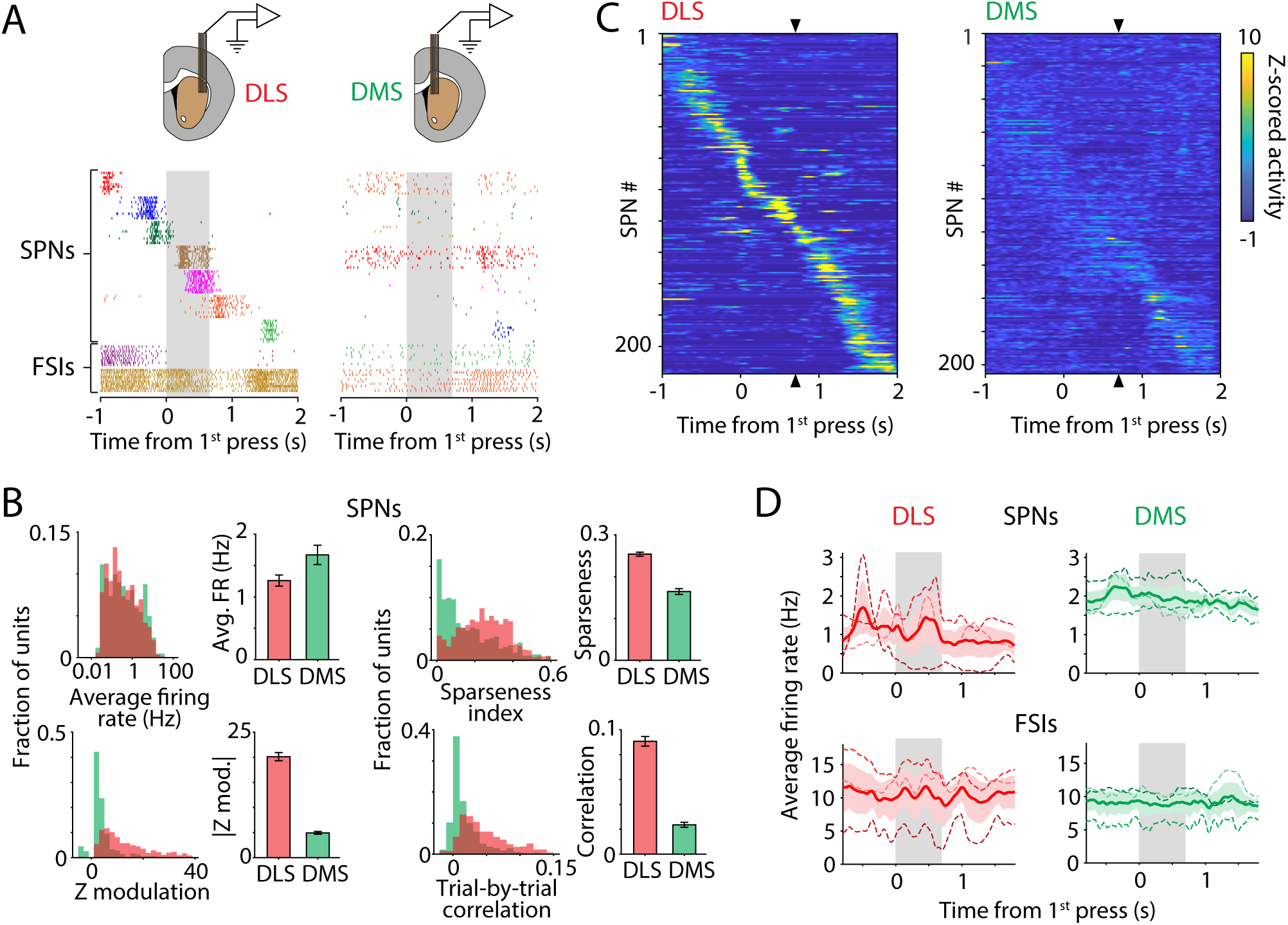
Units in DLS are significantly more modulated than units in DMS during the execution of a learned motor sequence. **A.** (Top) Schematic of multi-tetrode array recordings in the DLS (left) and the DMS (right) of animals performing the timed lever pressing task (Figure 1C). (Bottom) Raster plots showing spiking activity over 10 trials of 7 simultaneously recorded putative spiny projection neurons (SPNs) and 2 putative fast spiking interneurons (FSIs) from the DLS and DMS, aligned to the 1^st^ lever press in the timed lever pressing task. Grey shaded region indicates period between 1^st^ and 2^nd^ lever presses. **B.** Statistics of task-aligned activity, including average firing rate during the trial-period (top left), maximum modulation of Z-scored firing rate during the trial-period (bottom left), sparseness (top right) and average trial-to-trial correlation of task-aligned spiking (bottom right) in putative spiny projection neurons (SPNs) recorded in the DLS (red, n=819 units from 3 rats) and DMS (green, n=422 units from 3 rats). Bars and error-bars represent mean and SEM, respectively, across units. In this and all subsequent figures, we only included units with a minimum firing rate of 0.25 Hz in the task. **C.** Peri-event time histograms (PETHs) of Z-scored activity of SPNs recorded in the DLS (left) and DMS (right) of example rats. Units have been sorted by the time of their peak activity, in a cross-validated manner. The sorting index was calculated from PETHs generated using half the available trials for each unit, and then used to sort PETHs generated using the remaining trials. Triangles indicate time of the second lever press. **D.** Average firing rate over all SPNs (top) and FSIs (bottom) recorded in the DLS (left, red, n=819 SPNs and 284 FSIs) and DMS (right, green, n=422 SPNs and 212 FSIs). Individual rats are indicated by shaded dashed lines (n=3 for each group) and the average across rats by the solid line. Red and green shading represents SEM. Grey shaded region represents the time between the first and second lever press in the motor sequence.

Although DLS and DMS units had similar average firing rates during the task (Figure 2B), we found that spiking in DLS units was modulated to a far greater extent (Figure 2B). Their task-aligned activity was also more similar across trials (Figure 2B). Spiny projection neurons (SPNs) in the DLS also had much sparser activity patterns, often spiking only at one specific time during the behavior (Figure 2B). In contrast, SPNs in the DMS had more distributed activity patterns, resembling those of fast spiking interneurons (FSIs) in both striatal regions (Supplementary Figure 2A).

### DLS is continuously active throughout the learned motor sequence

That unit activity in the DLS is significantly modulated during the execution of the learned motor sequences is largely in agreement with previous reports on the involvement of the DLS, but not the DMS, in over-trained behaviors^25, 57^ (but see reference^58^). However, the way in which population activity in the DLS varies over the time-course of the behavior differs markedly across paradigms^13, 22, 23, 59^, a difference which has inspired the two major models of BG function (Figure 1A).

Results supporting the ‘action selection’ model show that neurons in the DLS are preferentially active at the beginning and end of over-trained motor sequences^22–24^. We note that the repetitive behaviors used in these tasks – locomotion and simple lever pressing – can be executed without BG involvement^13, 58, 60, 61^, likely by mid-brain and brainstem motor controllers.

Studies that train animals to modify the vigor of established behaviors towards a performance goal, such as in a timing task, have observed a more continuous representation of the learned behavior in the DLS^13^. This has been interpreted as the DLS activity modulating the vigor of ongoing motor patterns, such as locomotion^7, 45^.

To address the degree to which the neural representation of the learned motor sequences we train conform to either of these models, we first examined how activity in DLS neurons is distributed over the length of the motor sequence (Figure 2C). We found that the average activity in the DLS did not resemble a start/stop-like representation; instead, both SPNs and FSIs were continuously active throughout the motor sequence (Figure 2D). For individual animals, we found the distribution of average unit activity to be non-uniform and idiosyncratic (Figure 2D), reflecting the individually distinct behavioral solutions our trial-and-error learning paradigm produces^31^. Based on these results, we conclude that DLS SPNs represent the learned motor sequences in a sparse manner at the level of single neurons, and in a continuous manner across the population.

### DLS encodes low-level details of a learned motor sequence

Our results were not well explained by the ‘action selection’ model, according to which we would have expected DLS activity to bracket the over-trained behavior at its start and end^22, 23^. On the other hand, the continuous representation we observe in the DLS may be consistent with the “vigor” model, which proposes that the BG regulates the vigor of ongoing movements^7, 10, 13^. If this were the case, we would expect activity in the DLS to reflect vigor-related kinematic variables, such as movement speed^12, 13^.

Alternatively, since the motor cortex-independent motor sequences we train are likely controlled by brainstem and mid-brain circuits, they may rely on the BG in a way, and to an extent, previously not fully appreciated. If the BG indeed play the role of a controller, we would expect the DLS to encode additional details about the motor sequence, such as the direction or timing of its constituent movements.

To determine whether the activity we observed in the DLS is consistent with either of these models, we used a generalized linear model framework to probe which task-related parameters were encoded in the activity of individual DLS units (Figure 3A). As a control, we also determined the extent to which these parameters were encoded in the activity of DMS units, which were far less task-modulated (Figure 2B).

**Figure 3:**
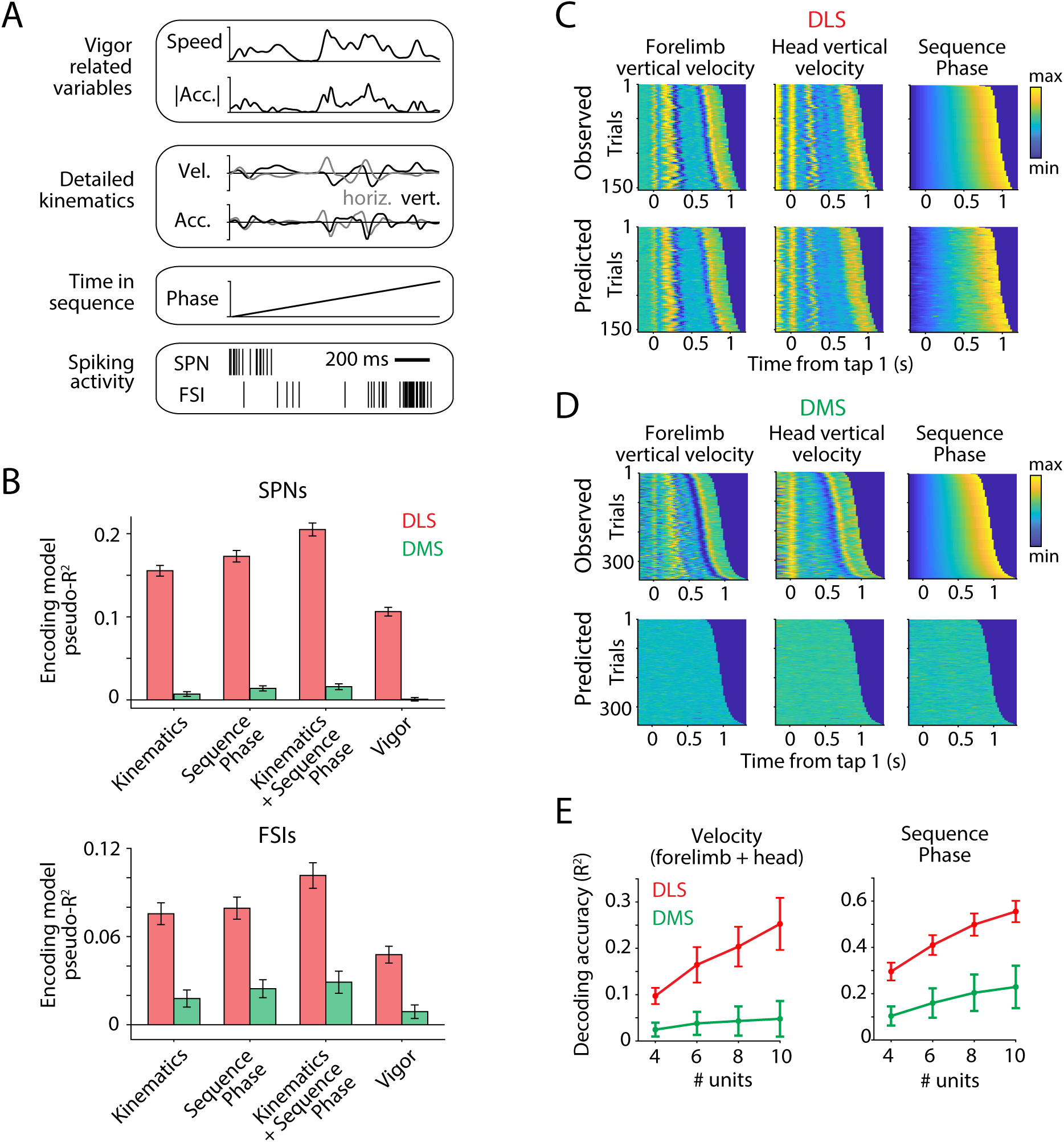
DLS, but not DMS, represents execution-level details of a learned motor sequence. A-B. Encoding analyses. **A.** Schematic of encoding analysis. We used generalized linear models to measure how well behavioral features such as vigor related variables (speed and magnitude of acceleration of both forelimbs and the head, top), movement kinematics (horizontal and vertical components of velocity and acceleration of both forelimbs and the head, middle), and inferred phase of the motor sequence (linearly scaled time between the first and second lever presses, bottom) predicted moment-by-moment changes in the spiking of striatal SPN and FSI units. Only kinematic data from the contralateral forelimb is shown for clarity. **B.** Goodness of fit, measured using the cross-validated pseudo-R^2^ (see Methods), of encoding models that use detailed information about the motor sequence, such as its kinematics and sequence phase, or vigor-related variables, to predict the instantaneous activity of putative SPNs (top) and FSIs (bottom) recorded in the DLS (red, n=474 SPNs and 237 FSIs from 3 rats) and DMS (green, n=209 SPNs and 142 FSIs from 2 rats). Bars and error-bars represent mean and SEM, respectively, across units. Only neurons that had a minimum firing rate of 0.25 Hz in the task were included in this analysis. **C-E.** Decoding analyses. **C.** (Top) Measurements of vertical velocity of the contralateral forelimb (left) and head (middle), and inferred sequence phase (right) for all trials in a representative task session of a DLS-implanted rat. Trials are aligned to the first lever press and sorted by the inter-press interval. (Bottom) Cross-validated predictions of vertical velocity and sequence phase by a neural network decoder that takes as input the instantaneous activity of all simultaneously recorded neurons. **D.** Same as in **C** but for a representative task session of a DMS-implanted rat. **E.** Average accuracy with which instantaneous velocity (left) and sequence phase (right) can be decoded by a neural network decoder as a function of number of units in the DLS (red) and DMS (green). Decoding accuracy was quantified by the fraction of variance explained (R^2^) between observed and predicted data, averaged over 4 cross-validation folds. We decoded both horizontal and vertical components of velocity of both forelimbs and the head. Ensembles of neurons of the indicated sizes (4 to 10) were generated by subsampling from sessions in which we had recorded at least 10 neurons simultaneously. Decoding accuracy was averaged over all such sessions for individual rats and then across rats in each group (n=3 for DLS and n=2 for DMS). Error-bars represent SEM across rats. Only neurons that had a minimum firing rate of 0.25 Hz in the task were included in this analysis.

We found that the details of how learned motor sequences are executed, such as the velocity and acceleration of the forelimbs and head, and the time within the motor sequence (Figure 3B, Supplementary Figure 3A), explained the activity of individual DLS units far better than scalar, vigor-related variables such as speed and magnitude of acceleration (Figure 3B, Supplementary Figure 3A). In contrast, all movement-related variables were encoded to a much lesser extent in the activity of DMS units (Figure 3B).

Our results thus far suggest that the DLS, but not the DMS, encodes the details of the learned motor sequences, such as their kinematics and timing (Figure 3B). However, it is not clear how complete of a representation this is, and whether these parameters can be reliably decoded from populations of DLS units.

To address this, we used a multilayer neural network decoder to test the degree to which simultaneously recorded populations of DLS or DMS units can decode the instantaneous velocity (both horizontal and vertical components) of the rats’ forelimbs and head during the task or the time within the sequence (see Methods, Figure 3C). We found that instantaneous velocities and timing could be accurately decoded from the DLS (Figure 3C), with decoding accuracy improving with number of neurons (Figure 3E). However, this was not the case with the DMS (Figure 3D-E). Indeed, we could not decode details of the motor sequence to any significant degree even from ensembles of 10 simultaneously recorded DMS units (Figure 3E).

### Movement encoding in DLS is independent of motor cortex

Thus far we have found that neurons in the DLS encode execution-level details of the learned motor sequences. This means that DLS, beyond modulating the vigor of the behavior, has information to control its detailed kinematic structure. If the neural representation in DLS is indeed involved in generating the kinematic structure of behavior, it ought to be independent of motor cortex, which we know from a previous study is not necessary for executing the motor sequences we test^31^.

To probe the dependence of behaviorally-locked striatal dynamics on motor cortical input, we recorded from the DLS in expert animals in which motor cortex had been lesioned after training (Figure 4A, n=1435 units in total, n=3 rats). As we have reported previously, large bilateral motor cortex lesions in expert animals did not materially affect the learned behaviors^31^ (Figure 1D).

**Figure 4:**
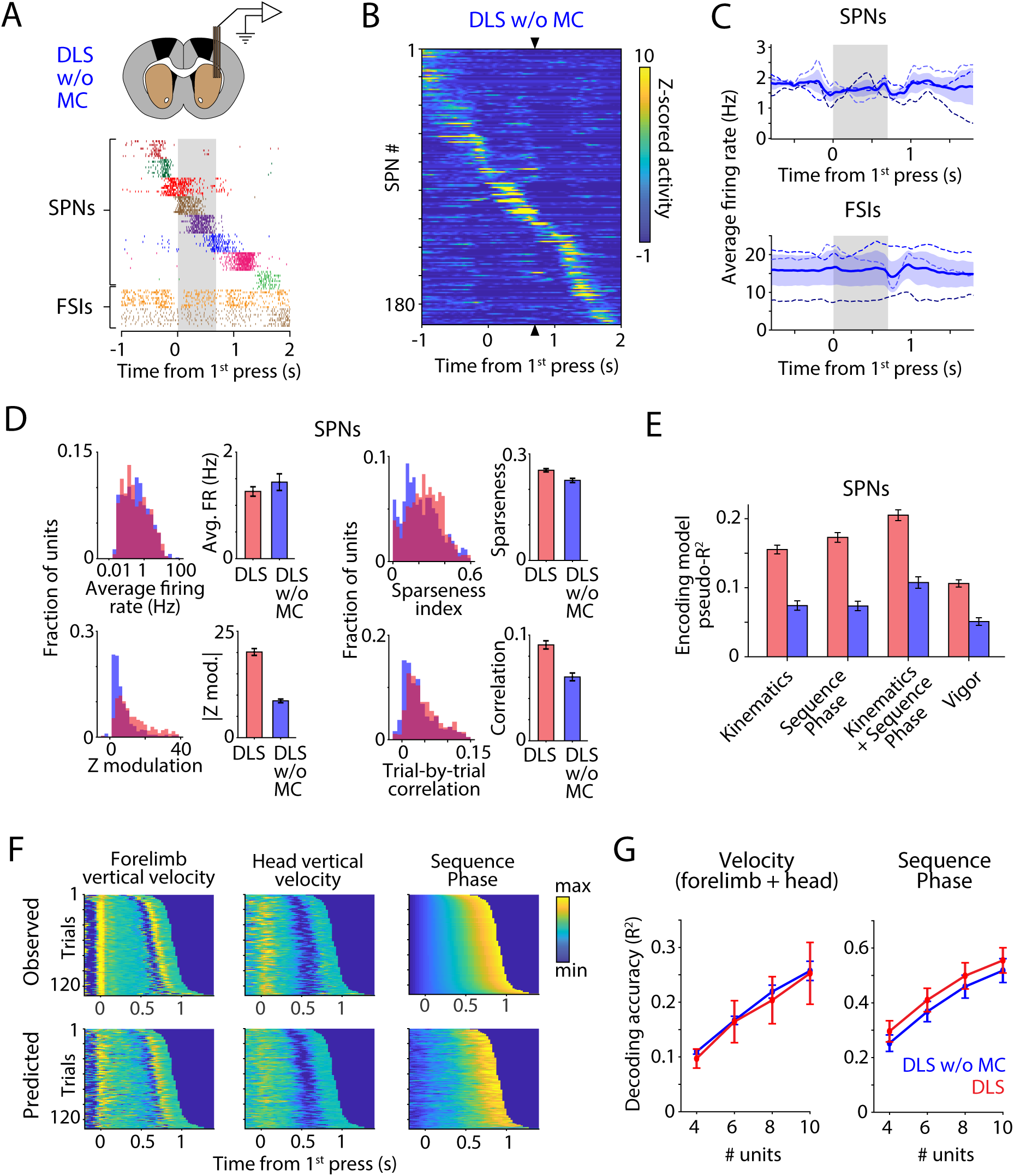
Movement encoding in DLS is independent of motor cortex. **A.** (Top) Schematic of multi-tetrode array recordings in the DLS of a motor cortex (MC)-lesioned rat. (Bottom) Raster plots showing spiking activity over 10 trials of 7 simultaneously recorded putative spiny projection neurons (SPNs) and 2 putative fast spiking interneurons (FSIs) from the DLS of a motor cortex-lesioned rat, aligned to the first lever press. Grey shaded region indicates period between 1^st^ and 2^nd^ lever presses. **B.** Peri-event time histograms (PETHs) of Z-scored activity of SPNs recorded in the DLS of an example motor cortex-lesioned rat. Units are sorted, in a cross-validated manner, by the time of their peak activity. The sorting index was calculated using PETHs generated from half the available trials for each unit, and then used to sort PETHs generated from the remaining trials. Triangles indicate time of the second lever press. **C.** Average firing rate over all SPNs (top, n=612 units) and FSIs (bottom, n=298 units) recorded in the DLS of motor cortex-lesioned rats. Individual rats are indicated by shaded dashed lines (n=3) and the average across rats by the solid line. Blue shading represents SEM. Grey shaded region represents the time between the first and second lever presses in the motor sequence. **D.** Statistics of task-aligned activity including average firing rate during the trial-period (top left), maximum modulation of Z-scored firing rate during the trial-period (bottom left), sparseness (top right) and average trial-to-trial correlation of task-aligned spiking (bottom right) of putative spiny projection neurons (SPNs) recorded in the DLS of intact (red, n=819 units from 3 rats) and motor cortex-lesioned (blue, n=612 units from 3 rats) rats. Bars and error-bars represent mean and SEM, respectively, across units. **E.** Goodness of fit, measured using the cross-validated pseudo-R^2^ (see Methods), of encoding models that use detailed information about the motor sequence such as its kinematics and sequence phase, or vigor-related variables, to predict the instantaneous activity of putative SPNs recorded in the DLS of intact (red, n=474 units) and motor cortex-lesioned (blue, n=292 units) rats. Bars and error-bars represent mean and SEM, respectively, across units. **F.** (Top) Measurements of vertical velocity of the contralateral forelimb (left) and head (middle), and inferred sequence phase (right) for all trials in a representative task session of a DLS-implanted motor cortex-lesioned rat. Trials are aligned to the first lever press and sorted by the inter-press interval. (Bottom) Cross-validated predictions of vertical velocity and sequence phase by a neural network decoder that takes as input the instantaneous activity of all simultaneously recorded neurons. **G.** Average accuracy with which instantaneous velocity (left) and sequence phase (right) can be decoded by a neural network decoder as a function of number of units in the DLS of intact (red) and motor cortex-lesioned (blue) rats. Decoding accuracy was quantified by the fraction of variance explained (R^2^) between observed and predicted data, averaged over 4 cross-validation folds. We decoded both horizontal and vertical components of the velocity of both forelimbs and the head. Ensembles of neurons of the indicated sizes (4 to 10) were generated by subsampling from sessions in which we had recorded at least 10 neurons simultaneously. Decoding accuracy was averaged over all such sessions for individual rats and then across rats in each group (n=3 for intact DLS and motor cortex-lesioned groups). Error-bars represent SEM across rats.

DLS units in motor cortex-lesioned rats had similar firing rates to those in intact rats and were also active over the entire duration of the motor sequence (Figure 4B-D). However, there were subtle but significant differences between the activity of DLS units in motor cortex-lesioned and intact rats. First, units in lesioned rats were less modulated during the task (Figure 4D, Supplementary Figure 2B), and their activity patterns were also less sparse and more variable from trial-to-trial (Figure 4D, Supplementary Figure 2B).

Furthermore, encoding analyses showed that the detailed kinematics and timing of the motor sequence were less effective at predicting the instantaneous activity of individual DLS units in motor cortex-lesioned animals as compared to those in intact animals. Note, however, that these features explained much more of the variance in unit activity than did vigor-related variables such as speed (Figure 4E, Supplementary Figure 2C, Supplementary Figure 3B). Since trial-to-trial variability in movement kinematics was similar for motor sequences performed by motor cortex-lesioned and intact rats (the average pairwise correlation of limb trajectories across trials was 0.70 ± 0.09 and 0.77 ± 0.09 for intact and motor cortex lesioned rats, respectively; mean ± SEM), this implies that removal of motor cortex led to an increase in neural variability in the striatum that is not reflected in, or originating from, the movements.

Whatever the source of the neural variability, if the BG do indeed play an essential role in controlling the details of the learned behavior, its population activity should still reflect kinematics and timing, despite this increase in variability. To probe this, we decoded instantaneous velocity from the spiking activity of DLS units in motor cortex-lesioned animals (Figure 4F-G). We found that decoding accuracy from ensembles of units was similar in lesioned and intact animals across a range of ensemble sizes (Figure 4G), consistent with the DLS having a similar amount of information about the execution-level details of the behavior with and without motor cortex.

These results suggest the sensorimotor arm of the BG represents kinematic structure of learned motor sequences and indicate that this pathway may function to influence activity in subcortical motor controllers to generate task-specific stereotyped motor sequences, independently of motor cortex.

### DLS is necessary for executing learned motor sequences

While our DLS recordings showed a continuous and detailed representation of the learned motor sequences, it remains an open question whether this activity is causal for controlling execution-level details of the learned behavior. Alternatively, it could be dispensable for the behavior altogether and simply reflect ongoing motor activity in essential non-motor cortical circuits from which DLS receives input^50, 53^. To address this directly, we lesioned the DLS bilaterally in expert animals (Methods; Figure 5A, Supplementary Figure 1B, n=7 rats) and investigated whether and how the loss of DLS activity affects the execution of the learned behaviors. For comparison, we lesioned the DMS (Figure 5A, Supplementary Figure 1B, n=5 rats), whose neurons are markedly less correlated with the kinematic details of the behaviors (Figure 3). To control for the injections and related surgery procedure, we also did control injections into DLS in a separate cohort of animals (Figure 5A, n=5 rats).

**Figure 5:**
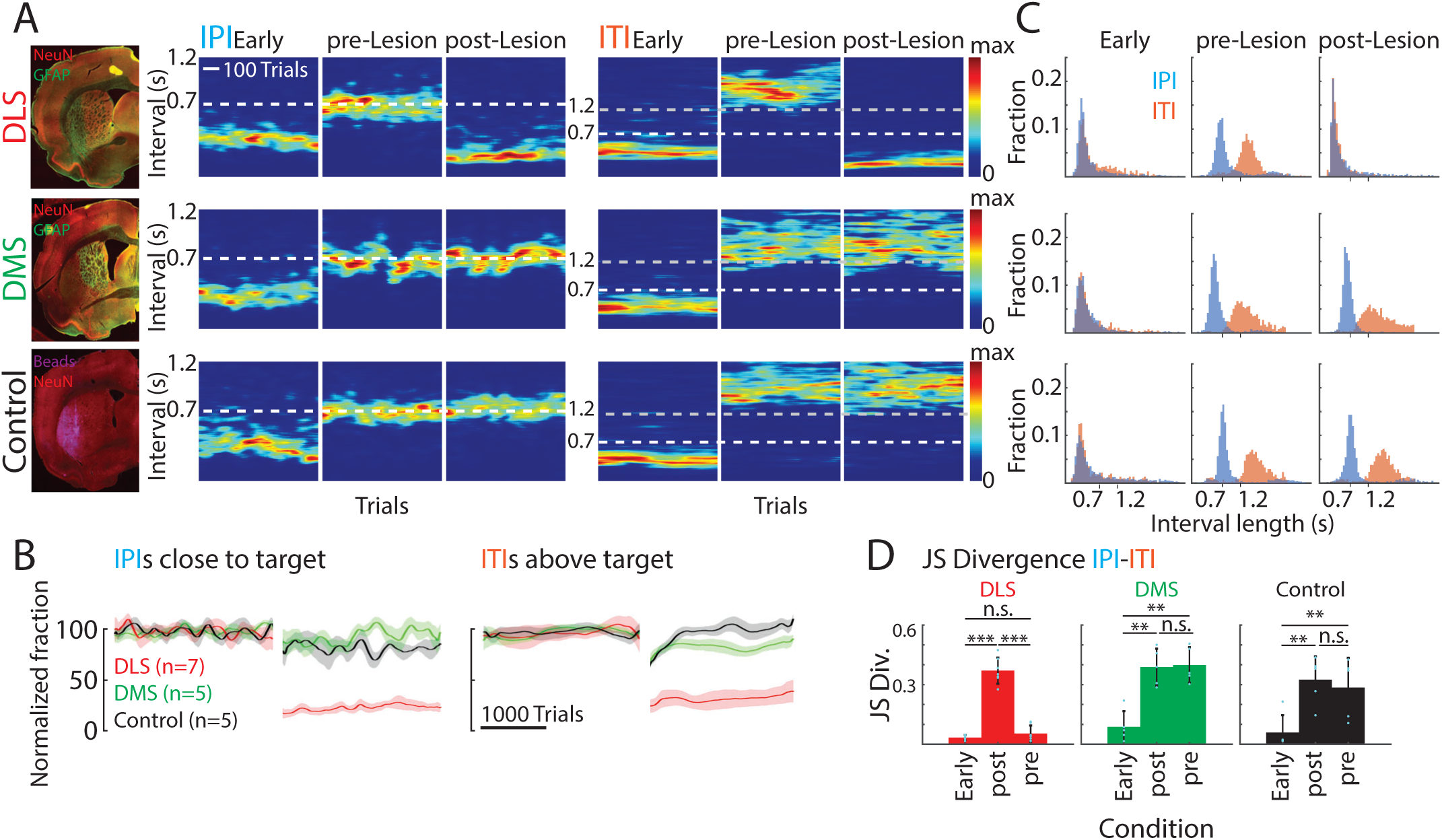
Lesions of DLS, but not DMS, degrade the performance in our timed lever pressing task. **A.** Representative examples of performance in the motor task (see Figure 1C) in animals subjected to different manipulations (DLS lesion, DMS lesion, DLS control injection). *Left*: Histological images of the manipulations. *Right*: Heatmaps of the probability distributions of the IPIs and ITIs for the example animals early in training, and before and after the manipulations. **B.** Average performance across animals (DLS n=7 rats; DMS n=5, Control n=5) for manipulations as in **A**, normalized to performance before the manipulation. Left: Fraction of trials with IPIs close to the target (700 ms +/−20%). Right: Fraction of trials with ITIs above the threshold of 1.2 s. **C.** Distributions of interval lengths between taps for the animals shown in **A** early in training, and before and after the manipulations. **D.** Dissimilarity between the IPI and ITI distributions across animals early in training, and before and after the manipulations. Shown is the JS Divergence as a measure of dissimilarity. The reduction of the JS Divergence after DLS lesion indicates an increase in the overlap of the two interval distributions to levels observed early in training. Error bars represent standard error of the mean (SEM). \*\**P* < 0.01, \*\*\**P* < 0.001.

Lesions of the DLS drastically impaired the animals’ performance. While rats were still actively engaged in the task, their IPIs decreased relative to pre-lesion and they became, on average, more variable (Figure 5A, Supplementary Figure 4A). This, in turn, led to a significant drop in the number of ‘successful’ trials, defined here as the IPI being within 20% of the target (700 ms, Figure 5B). Notably, post-lesion performance was indistinguishable from the early stages of learning (Figure 5A, Supplementary Figure 4A), and did not recover even after extended periods of additional training (Supplementary Figure 4A), suggesting that DLS is also required for relearning the task. In contrast, lesions of the DMS did not affect the performance of expert animals beyond what can be expected after control injections into DLS and subsequent recovery (Figure 5A-B, Supplementary Figure 4A).

In addition to mastering the prescribed IPI target (700 ms), normal animals also learn to withhold lever pressing after unsuccessful trials for at least 1.2 seconds (the inter-trial interval, ITI) - a requirement to initiate a new trial (Figure 1C). As animals learn the structure of the task during training, they develop separate strategies for timing the two intervals (Figure 5A,C), as evidenced by distinct peaks in the overall lever press interval distributions (Figure 5C). After DLS lesion, however, the mean ITI duration is not only reduced (Figure 5A-B; Supplementary Figure 4A), but the distinction between the IPIs and ITIs is completely lost (Figure 5C-D). Interestingly, the temporal structure of the animals’ lever pressing behavior reverts to what is seen in early stages of training (Figure 5C-D). Thus, in contrast to DMS lesioned animals and animals subject to control injections, DLS lesioned animals were unable to reproduce or relearn the previously acquired task structure. (Figure 5).

It has been proposed that motor deficits in striatum-related disorders, like Parkinson’s disease (PD), are due not to the loss of striatal function, but rather to altered dynamics in striatum causing the BG to produce aberrant output^62–65^. In support of this idea, lesions of the internal segment of the globus pallidus internal segment (GPi), one of the main output nuclei of the BG, have proven an effective treatment for dyskinesias in PD^21, 66, 67^. Thus, impairments observed after DLS lesions might either be due to loss of instructive DLS activity, or, alternatively, to the production of aberrant BG dynamics that disrupts the task-related activity of downstream control areas^10^. To distinguish between these possibilities, we lesioned the rat homolog of the GPi, the endopenduncular nucleus (EP), in an additional group of animals (Supplementary Figure 5A, n=5 rats). This manipulation affected task performance in a similar way to DLS lesions (Supplementary Figure 5A-D). Taken together, these results show that the DLS are required for producing the learned motor sequences we train.

### DLS lesions disrupt the learned motor sequences

While we have shown that DLS lesions impair task performance, distinguishing between different models of BG function would be helped by describing the specific motor deficits in more details. On the one hand, performance could suffer from changes to the speed or amplitude of the learned motor sequences - deficits consistent with the “vigor” model^10, 11, 68^. On the other hand, deficits could result from an inability to generate the learned movement sequences altogether, an outcome that would suggest that the BG are actively orchestrating motor controllers in downstream circuits^3, 32, 34, 35, 37, 69^. To better arbitrate between these possibilities, we used video-based motion tracking^55, 56^ to compare the detailed kinematics of task-associated movement patterns before and after bilateral DLS and DMS lesions (see Methods).

We have previously shown that rats ‘solve’ the timed lever press task by consolidating highly idiosyncratic and stereotyped motor sequences^31^. While the trial-and-error learning process and the kinematically rather unconstrained nature of the task lead to idiosyncratic and diverse behavioral solutions, once a rat converges on a successful motor sequence, it tends to reproduce it in the very same form over long periods of time and also across weeks of rest^31^, suggesting the formation of stable task-specific motor memories.

In line with our analyses of performance metrics, we found that the learned motor sequences were faithfully reproduced after DMS lesions, suggesting that the associative arm of the basal ganglia does not contribute meaningfully to the execution of the learned motor skills. (Figure 6B,D). In contrast, movement patterns of DLS-lesioned rats changed dramatically (Figure 6A). While still fairly stereotyped, they did not at all resemble the pre-lesion motor sequences (Figure 6A). Instead of the highly idiosyncratic task-specific movement patterns characteristic of normal animals, the behaviors expressed by different animals after DLS lesion were surprisingly similar to each other (Figure 6C).

**Figure 6:**
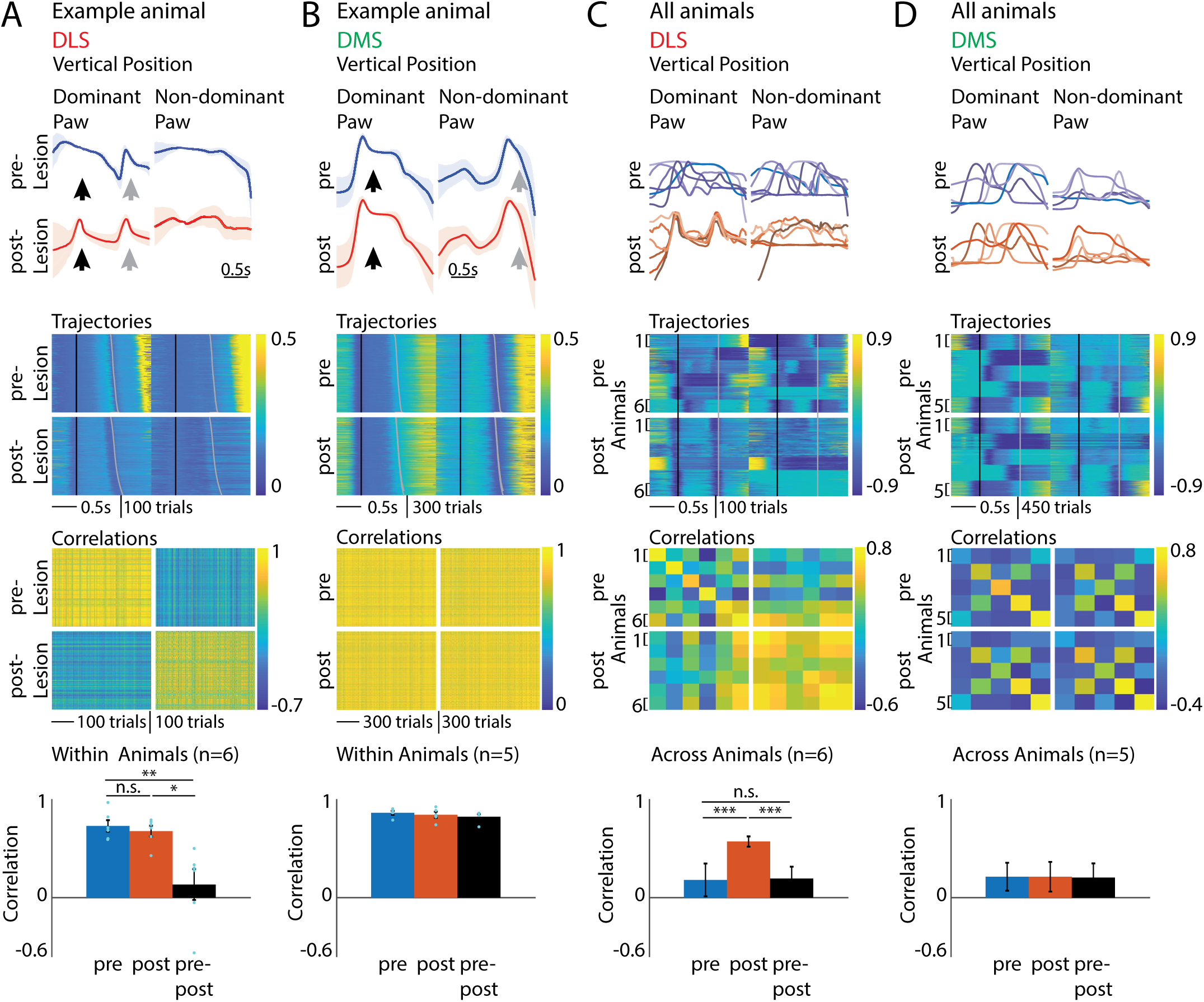
Lesions of the DLS, but not the DMS, lead to a loss of the idiosyncratic learned motor sequences and a regression to movement patterns common across animals. **A.** Within animal comparison of the forelimb trajectories associated with the task before and after DLS lesions. *Row 1*: Example average forelimb trajectories (position in the vertical dimension) before and after DLS lesion (calculated from trials close to the mean IPI (mean IPI +/−30 ms)). Black arrows indicate the 1^st^ press, grey arrows the 2^nd^ press. The forelimb performing the first lever press is regarded as the dominant one. *Row 2*: Vertical forelimb displacement in individual trials before and after DLS lesion for both limbs, sorted by IPI. Black lines mark the time of the 1^st^ press, grey lines of the 2^nd^. *Row 3*: Pairwise correlations of the forelimb trajectories shown above after linear time-warping of the trajectories to a common time-base (see Methods). *Row 4*: Average pairwise correlations within animals averaged across animals (n=6 rats). Mean +/−SEM. \**P* < 0.05, \*\**P* < 0.01. **B.** Like **A**, but for DMS lesions (n=5 rats). **C.** Comparison of forelimb movement trajectories across animals before and after DLS lesion. *Column 1:* Comparison of average trajectories (vertical displacement) of all animals before and after DLS lesion (n=6 rats). *Column 2:* Vertical forelimb displacement in randomly selected trials (80 per animal) of all animals before and after DLS lesion for dominant and non-dominant forelimbs, sorted by IPI. Black lines mark the time of the 1^st^ lever press, grey lines of the 2^nd^ press. *Column 3:* Pairwise correlations between the trials shown in column 2, averaged per animal. *Column 4:* Averages of the correlations shown in G) by condition (average of all pre-to-pre, post-to-post and pre-to-post correlations). Mean +/−SEM. \*\*\**P* < 0.001. **D.** Like **C**, but for DMS lesions (n=5 rats).

These results are not explained by DLS lesions affecting performance simply by altering the vigor of established motor sequences; rather, the complete loss of the learned motor sequences suggest an essential control function for the BG.

### DLS lesions cause a reversion to a species-typical lever pressing behavior

To better understand the nature of the post-lesion deficits and what they can tell us about BG’s control function, we analyzed the forelimb movement trajectories associated with individual lever presses. In expert animals these are highly idiosyncratic and also distinct for the first and second lever press in a sequence. Following DLS lesions, however, there was barely any distinction in how individual rats executed the first and second lever press (Figure 7A). The high similarity between the post-lesion lever presses within animals (Figure 7A), and between the full post-lesion trajectories (Figure 6C) across individual animals, prompted us to also compare the lever press movements across animals. In contrast to the idiosyncratic lever presses of intact animals, the forelimb trajectories of all lever presses after DLS lesions, both across first and second presses and across animals, were highly similar (Figure 7B) - a dramatic change from before the lesions when they were all largely distinct (Figure 7B).

**Figure 7:**
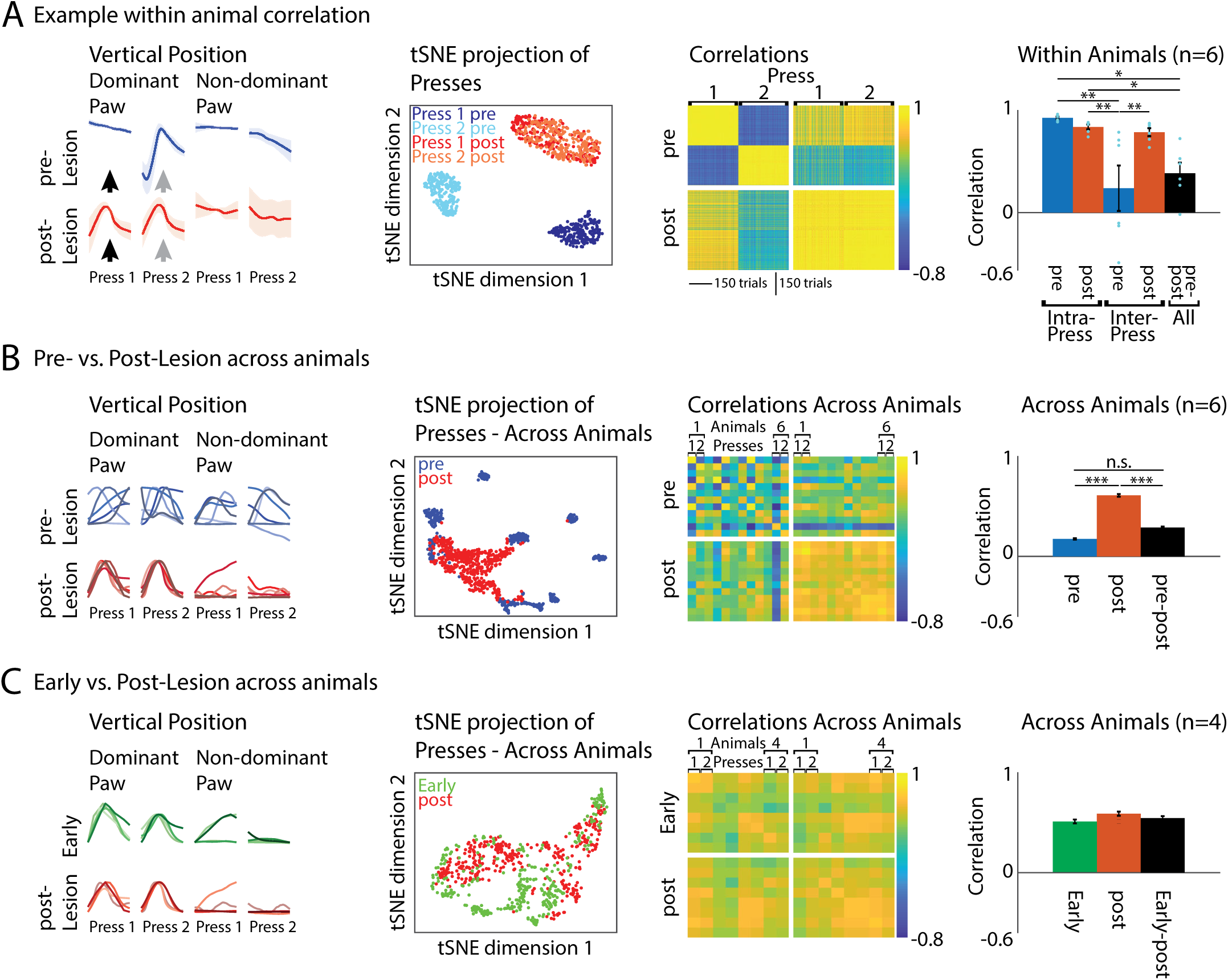
Lesions of the DLS lead to a loss of the idiosyncratic learned lever press movements and a regression to a press movement common across animals and similar to what is seen early in learning. **A.** Comparison of 1^st^ and 2^nd^ lever presses within animals before and after DLS lesion. *Column 1*: Average movement trajectories for the 1^st^ and 2^nd^ press before and after DLS lesion in an example animal. Black and grey arrows indicate the time of the 1^st^ and 2^nd^ press, respectively. *Column 2*: t-SNE projection of 1^st^ and 2^nd^ lever press trajectories before and after DLS lesion for the example animal from column 1. *Column 3*: Pairwise correlations between lever press associated forelimb movements of the example animal before and after DLS lesion. *Column 4*: Averages of within animal correlations across animals (n=6 rats) and conditions. Shown are correlations between forelimb trajectories before and after DLS lesion between the same presses (Intra-Press: 1^st^ to 1^st^ and 2^nd^ to 2^nd^) and between different presses (Inter-Press: 1^st^ to 2^nd^). Also shown are correlations between the before and after lesion conditions (All) across all 1^st^ and 2^nd^ presses. Mean +/−SEM. **B.** Comparison of 1^st^ and 2^nd^ lever presses across animals before and after DLS lesions. *Column 1*: Forelimb movement trajectories for the 1^st^ and 2^nd^ press before and after DLS lesion, overlaid for all tracked animals (n=6 rats). *Column 2*: t-SNE projection of all 1^st^ and 2^nd^ press trajectories before (blue) and after (red) DLS lesion for all animals from column 1. *Column 3*: Pairwise correlations between press trajectories of all animals before and after DLS lesion. Shown are average trial-to-trial correlations across individual presses (animal 1 press 1, animal 1 press 2, etc.). *Column 4*: Averages of across animal correlations between conditions. Shown are correlations between all presses before and all presses after DLS lesion. Also shown are correlations between all presses before and all presses after lesion. Mean +/−SEM. **C.** Comparison of 1^st^ and 2^nd^ lever presses across animals early in training and after DLS lesion. *Column 1*: Forelimb movement trajectories for the 1^st^ and 2^nd^ presses early in training and after DLS lesion (replotted from panel **B**), overlaid for all tracked animals (n=4 rats, subsample of rats in **B**, for which trajectories were available early in training). *Column 2*: t-SNE projection of all 1^st^ and 2^nd^ press trajectories early in training and after DLS lesion for all animals from column 1. *Column 3*: Pairwise correlations between press trajectories of all animals early in training and after DLS lesion. Shown are average trial-to-trial correlations across individual presses (animal 1 press 1, animal 1 press 2, etc.). *Column 4*: Averages of across animal correlations between conditions. Shown are correlations between all presses early and all presses after DLS lesion. Also shown are correlations between all presses early and all presses after lesion. Mean +/−SEM.

The high similarity in how DLS lesioned animals pressed the lever (Figure 7B), and the fact that performance decreases to levels seen early in training (Figure 5; Supplementary Figure 3A) led us to speculate that the DLS lesioned animals revert to a basal ganglia-independent species-typical lever pressing strategy, perhaps produced by control circuits in the brainstem^38, 70, 71^.

If rats indeed have an innate and favored means of pressing the lever, we argued that they would use it early in learning as a substrate for the trial-and-error learning process that follows. To test whether the post-lesion lever press movements resemble those seen early in learning, we compared the forelimb trajectories associated with lever presses for a subset of animals across these conditions (Figure 7C). Indeed, we found that the post-lesion and early pre-lesion lever press movements were highly similar across all animals (Figure 7C).

Overall, results from our lesion experiments show that activity in the DLS is necessary to produce the learned idiosyncratic motor sequences in our task. Lesions of the DLS completely disrupted the learned sequences, causing animals to regress to a species-typical, DLS-independent, motor pattern which allowed them to continue pressing the lever and collect reward.

## Discussion

We set out to test the role of the BG in the execution of motor skills, specifically the stereotyped task-specific motor sequences acquired in our task (Figure 1C,D). Our experimental results are not adequately explained by either of the established models of BG function, which posit that the BG are involved in selecting actions or modulating their vigor. Rather, our results point to an essential control function for the sensorimotor arm of the BG. The evidence came both from neural recordings, showing that DLS neurons encode the temporal structure and kinematics of the learned behavior (Figures 2,3), and from lesions of DLS (Figure 5,6) and the GPi (Supplementary Figure 5), which resulted in a complete loss of the learned motor sequences, and a reversion back to a species-typical task-related behavior (Figure 7). Interestingly, the task-relevant neural representations in the striatum were independent of motor cortex (Figure 4), as is the learned behavior itself^31^ (Figure 1D). Overall, our results suggest that the BG can function to control lower-level motor circuits to generate task-specific skilled behaviors.

### Relation to prior work on the BG and motor skill execution

At first, our findings may seem at odds with prior studies, which have come to different conclusions regarding BG’s function in motor skill execution. There is indeed a heterogeneity of results and conclusions to consider, even if we limit ourselves to studies in which rodents are tasked with pressing a lever or joystick^12, 23, 24, 58^. In most cases experimental outcomes have conformed to one of the established models of BG function - the ‘action selection’ or the ‘vigor’ hypotheses^12, 15, 23, 24, 58, 72^. Our results, in contrast, were not adequately explained by either of these models.

A constructive way to deal with the seeming discrepancies is to consider how the various studies differ in terms of the challenges posed by the behavioral tasks. For instance, several studies ascribing an ‘action selection’ role to the BG reward animals for producing kinematically and temporally unconstrained repetitive lever pressing behaviors^23, 24, 27^. Repetitive lever pressing itself is likely ‘solved’ by mid-brain/brainstem/spinal motor circuits and does not seem to require the BG^58, 60, 61^. This is further supported by our study, in which DLS lesioned rats, incapable of executing the previously acquired sequences, were perfectly capable of repetitive lever pressing (Figure 7). In these less constrained lever pressing tasks, therefore, BG’s main role may be to initiate and terminate activity in downstream circuits that control repetitive lever pressing (Figure 1A).

Other studies in which the results conformed better to the ‘vigor’ hypothesis (Figure 1A), differ in that they require animals to modulate the speed of movements or action sequences to successfully meet a performance criterion and obtain reward^12, 13, 15^. The more continuous neural representation in striatum seen during such behaviors is consistent with a role for the BG in modulating the overall vigor of ongoing movements and actions^7^.

In contrast to the aforementioned studies, training in our timed lever pressing task gradually changes the kinematic structure of the animal’s lever pressing movements, and also adds extraneous movement to time the prescribed delay between the presses^31^ (Figure 1D). Thus, what initially starts out as a species-typical repetitive lever pressing behavior (Figures 1D,7C), is shaped through training to become an idiosyncratic motor sequence quite distinct from the initial behavior (Figures 1D,6C,7B).

That the idiosyncratic motor sequences that emerge from training in our timed lever pressing task are qualitatively different from the behaviors in the oft-used repetitive lever pressing tasks, and that the BG ‘treat’ these tasks differently, is evident also when comparing the neural representations in the DLS and DMS^58^ across the tasks. For example, over-trained repetitive lever pressing behaviors appear to be encoded in a very similar manner in both the DLS and the DMS^58^, wherein SPNs preferentially encode the start and stop of the sequence^23, 58^. This lack of a more detailed representation in both dorsal striatal sub-regions is consistent with neither area being required for the control of the behavior^58, 60^. While acute inactivations of either DLS or DMS can lead to some subtle alterations in behavioral vigor, they do not materially affect the rats’ ability to perform lever press sequences^58^. In contrast, we see markedly different neural representations in the DLS and DMS during the execution of the skilled motor sequences we train (Figure 2), reflecting the differential roles these striatal sub-regions play in controlling the behavior (Figure 5).

Our study highlights the importance of carefully considering the specifics of the behavioral task when interpreting the results of a study. Differences in task designs across studies can add great value and serve to improve our understanding of the roles neural systems play in specific processes by allowing general hypotheses to be tested in different ways. In this spirit, we do not see our results as contradicting or invalidating previous theories of BG function. Our results do not, however, support their generality or exclusivity.

Instead, our study favors a view of the BG in which they are a versatile and flexible contributor to motor skill execution. If the actions needed to collect maximum reward can be generated by dedicated control circuits in the motor cortex or brainstem, then BG can simply select them and modulate their vigor (the ‘jukebox’ analogy introduced above, Figure 1A). However, if the reliable execution of novel movement patterns is needed, BG can get in on the act of shaping and controlling the details of the behaviors through their output to various motor control circuits (the ‘modern day DJ’ analogy, Figure 1A).

### BG control of brainstem and mid-brain motor circuits

That the BG interact with brainstem and midbrain motor centers is hardly a new idea. Indeed, in most vertebrates, these nuclei are the BG’s main targets^4, 32^. This phylogenetically older BG-pathway is also important for mammals to generate innate sequential behaviors, such as grooming^42, 43^. Not unlike the motor sequences we train, grooming comprises complex and fairly stereotypical behaviors that aren’t contingent on motor cortex^42^. Similar to our findings, some details of the grooming sequences are encoded in the DLS^39, 73^ and grooming syntax is disrupted with focal lesions of the DLS^43^. Thus, the learned behaviors we probe may tap into the same control circuits and mechanisms that have been shaped over evolution to generate robust species-typical sequential behaviors. That there are common substrates and mechanisms for encoding and generating innate and learned motor sequences in the brain suggest that there may be less of a distinction in their neural control than commonly assumed^74^.

### The nature of the control signals in the DLS

Given that the neural activity patterns we observe in the DLS reflect BG’s control function, it raises the question of how these control signals are generated. Our results, showing that behaviorally relevant striatal dynamics are maintained after motor cortex lesions (Figure 4), show that this activity does not require input from motor cortex as is commonly assumed^75, 76^. Recent modeling studies have suggested that sequence-associated neural dynamics in DLS could arise from recurrent activity within the striatum itself^77^. Since the striatum is an inhibitory structure, this would require a permissive excitatory drive which, in our motor cortex lesion rats, could be provided by either of its remaining inputs, from the thalamus or somatosensory cortex respectively^50, 53^. Alternatively, dynamics in DLS could be driven directly by behaviorally-locked activity in its non-motor cortical inputs. Future experiments will be required to arbitrate between these possibilities.

### What aspects of behavior are the BG controlling?

While our study suggests that dynamics in BG circuits can control the detailed structure of learned behaviors, the specific nature of this putative control signal is not clear. While we can decode detailed kinematics and timing of stereotyped motor sequences quite well from the activity of a few neurons in the DLS (Figure 3), it does not necessarily mean that DLS controls the kinematics.

Another possibility is that the BG control the sequential structure and/or timing of the learned motor sequences^78, 79^. In support of a role for the BG in motor timing, patients with Parkinson’s and Huntington’s disorder, in whom basal ganglia function is compromised, show deficits in timing tasks^80, 81^. The idea that DLS generates the temporal structure of learned sequential behaviors is inspired by work in songbirds, where the premotor nucleus HVC is involved in generating the temporal structure of their song – another example of a learned and stereotyped motor sequence^82^. Intriguingly, the sparse and behavior-locked activity patterns we see in SPNs are highly reminiscent of recordings of HVC projection neurons during singing^83^. The ‘timing neurons’ in HVC are thought to trigger muscle-specific control units in a downstream area, RA^82^. Likewise, the activity patterns in the DLS could drive BG output neurons to trigger control circuits in the midbrain/brainstem (Figure 1A, bottom panel). Similar to DLS lesions in our rats, removal of HVC abolishes the bird’s capacity to generate learned songs, but spares the production of species-typical calls^84^.

Disambiguating whether a brain structure encodes timing and/or kinematics is difficult in the context of a stereotyped sequential behavior, because the two aspects are highly correlated (Figure 1C,D). Addressing the proposed control function of the DLS will require either more subtle manipulations, such as cooling^85^, or new behavioral tasks in which kinematics and temporal structure can be more readily dissociated^86^.

In summary, our study probed the function of the BG through the lens of a motor cortex-independent learned behavior with rich and idiosyncratic kinematic structure. Our results, which did not conform to established models of how the BG contribute to motor skill execution, extend our understanding of their function to include an important role in controlling the execution-level details of learned motor sequences. The specifics of how this function is implemented in neural circuitry remains to be elucidated.

**Supplementary Figure 1:**
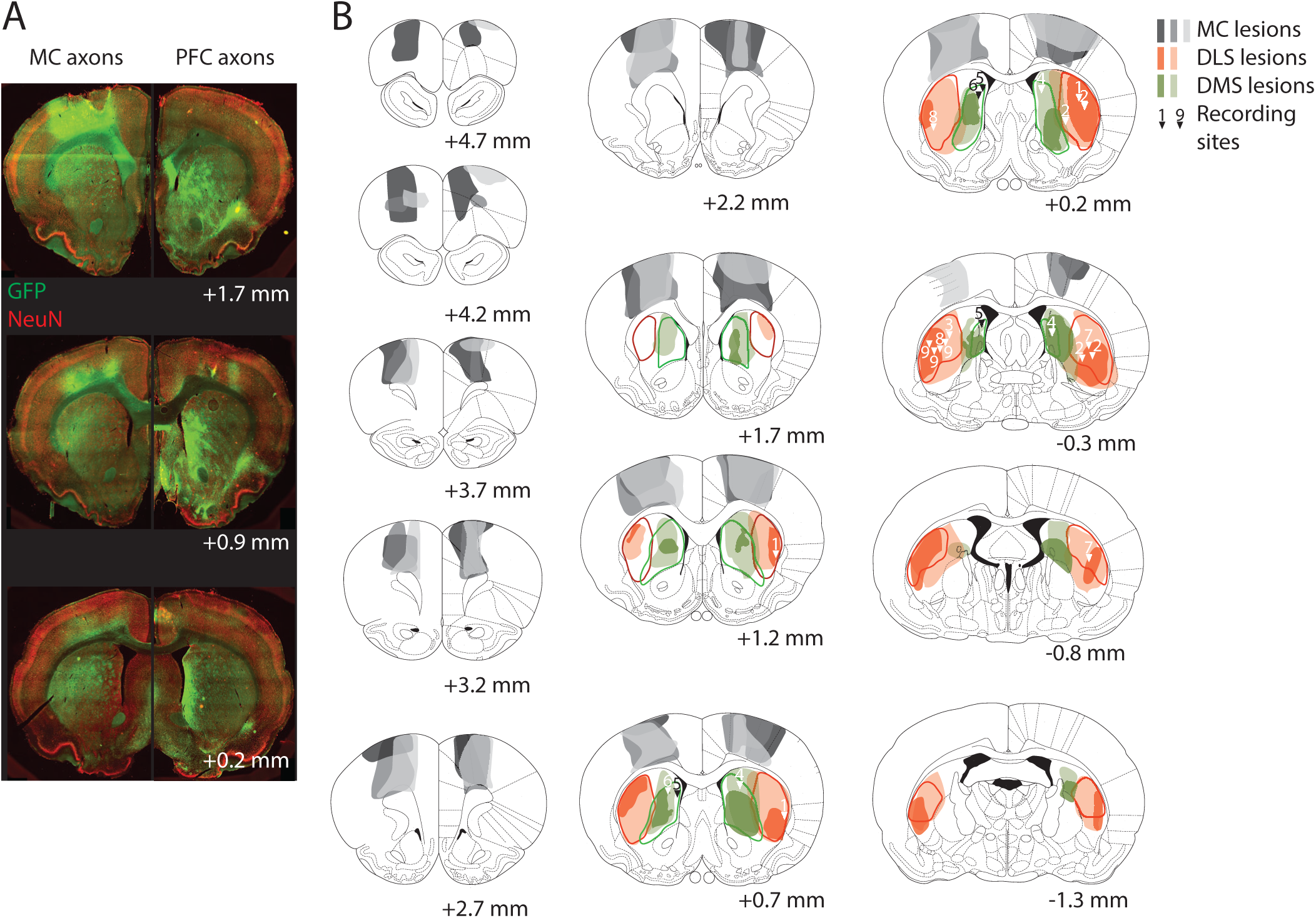
Striatal subdivisions, recording sites and extent of lesions. **A.** Virally-mediated fluorescent labeling of axons originating in either motor cortex (MC) or prefrontal cortex (PFC) to determine the outlines of the motor cortex-recipient dorsolateral striatum (DLS) and of the prefrontal cortex-recipient dorsomedial striatum (DMS), respectively. Based on the distinct projection patterns we estimated the extent of the DLS and DMS, respectively, along the anterior-posterior axis of the Striatum. **B.** DLS/DMS outlines, recording sites and lesion extents. The outlines of the DLS and DMS determined in **A** along the anterior-posterior axis are indicated by red and green lines, respectively. Locations of electrode implantation sites in DLS and DMS for electrophysiological recordings are marked with arrowheads. Numbers indicate different animals. For some animals several recording locations were determined, due to individual tetrode bundles of our recording arrays spreading apart during implantation. The extents of motor cortex lesions in three recorded animals are marked in different shades of grey for individual animals. The extents of the DLS and DMS lesions are marked as shaded red and green areas, respectively. Lighter areas indicate the extent of the largest lesion across animals at a given anterior-posterior position, darker areas indicate the extent of the smallest lesion.

**Supplementary Figure 2:**
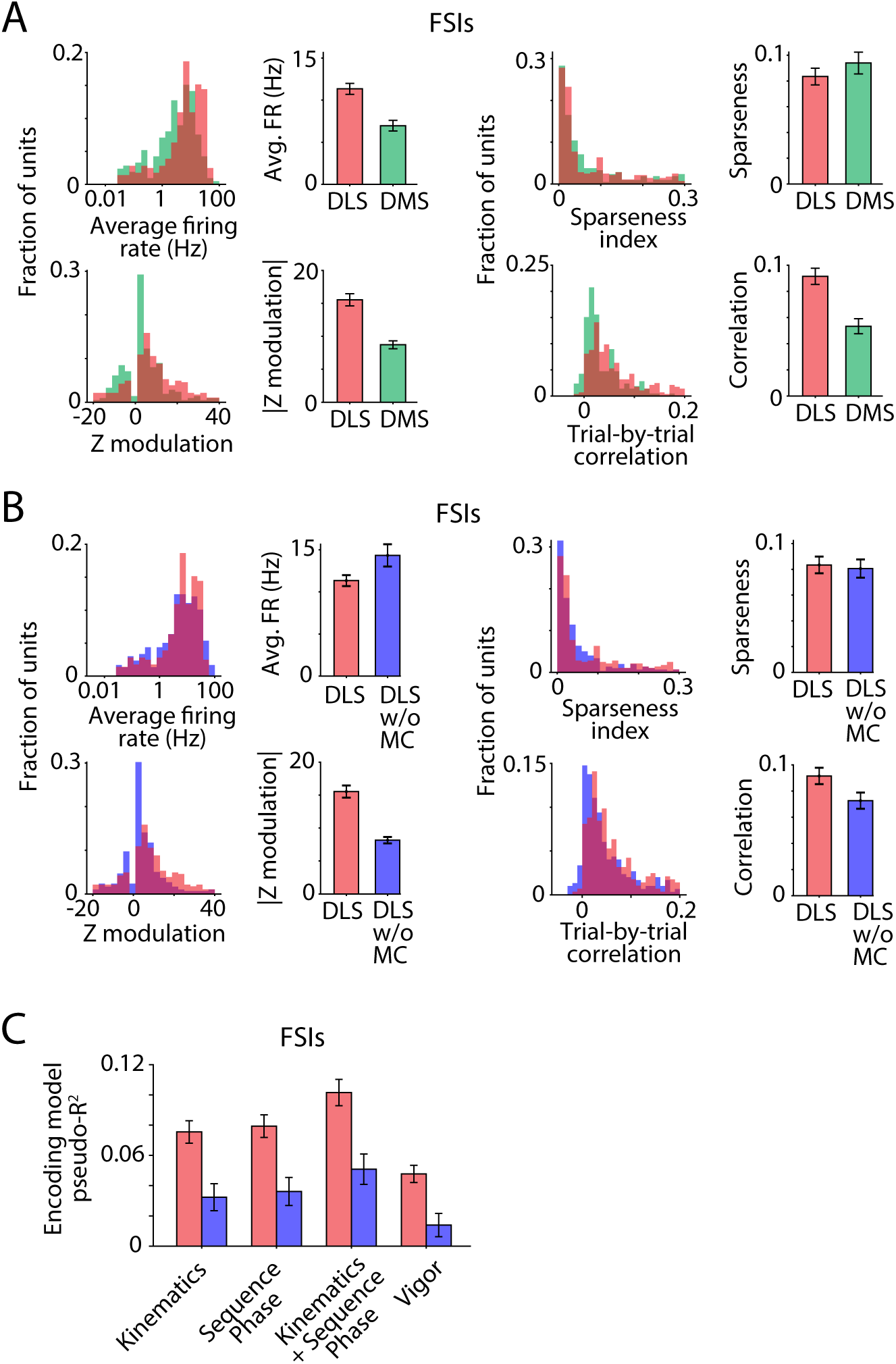
Task-aligned activity and encoding of movements by striatal FSIs. **A-B.** Statistics of task-aligned activity including average firing rate during the trial-period (top left), maximum modulation of Z-scored firing rate during the trial-period (bottom left), sparseness (top right) and average trial-to-trial correlation of task-aligned spiking (bottom right) of putative fast spiking interneurons (FSIs) recorded in the DMS (**A**, green, n=212 units), and DLS of intact (**A-B**, red, n=284 units) and motor cortex-lesioned (**B**, blue, n=298 units) rats. Bars and error-bars represent mean and SEM, respectively, across units. Only units with a minimum firing rate of 0.25 Hz in the task were included in this analysis. **C.** Goodness of fit, measured using the cross-validated pseudo-R^2^ (see Methods), of encoding models that use detailed information about the motor sequence such as its kinematics and sequence phase, or vigor-related variables, to predict the instantaneous activity of putative FSIs recorded in the DLS of intact (red, n=237 units) and motor cortex-lesioned (blue, n=209 units) rats. Bars and error-bars represent mean and SEM, respectively, across units.

**Supplementary Figure 3:**
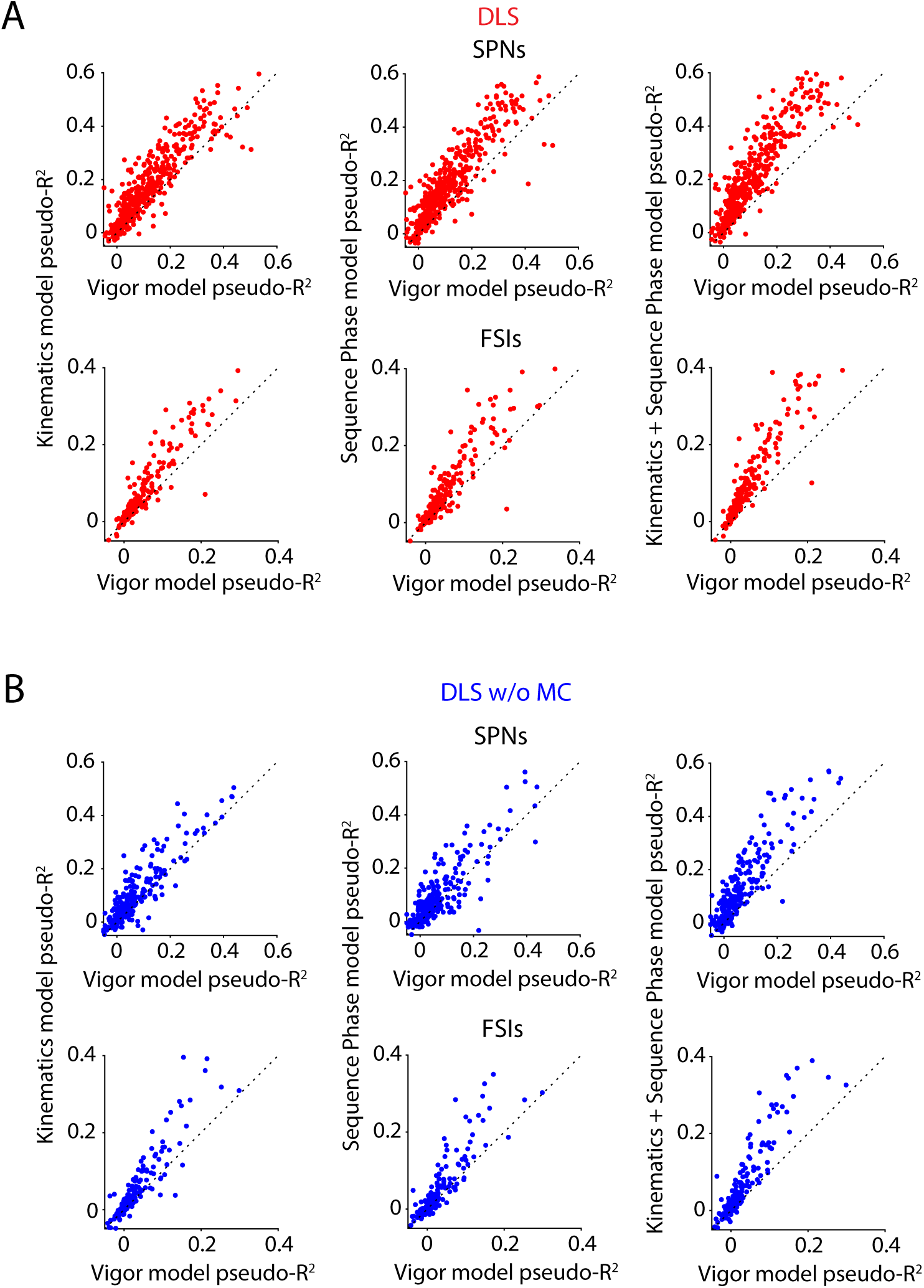
Comparison of encoding of detailed kinematic structure versus vigor by individual striatal neurons. **A-B.** Scatter plots comparing the goodness of fit, measured using the cross-validated pseudo-R^2^ (see Methods), between encoding models that use detailed information about the motor sequence such as its kinematics (left column) and sequence phase (middle column) or their combination (right column) and encoding models that use vigor-related variables for individual SPN (top) and FSI (bottom) units (indicated by points) recorded in the DLS of intact (**A**, blue, n=474 SPNs and 237 FSIs) and motor cortex-lesioned (**B**, red, n=292 SPNs and 209 FSIs) rats. Dashed line represents unity.

**Supplementary Figure 4:**
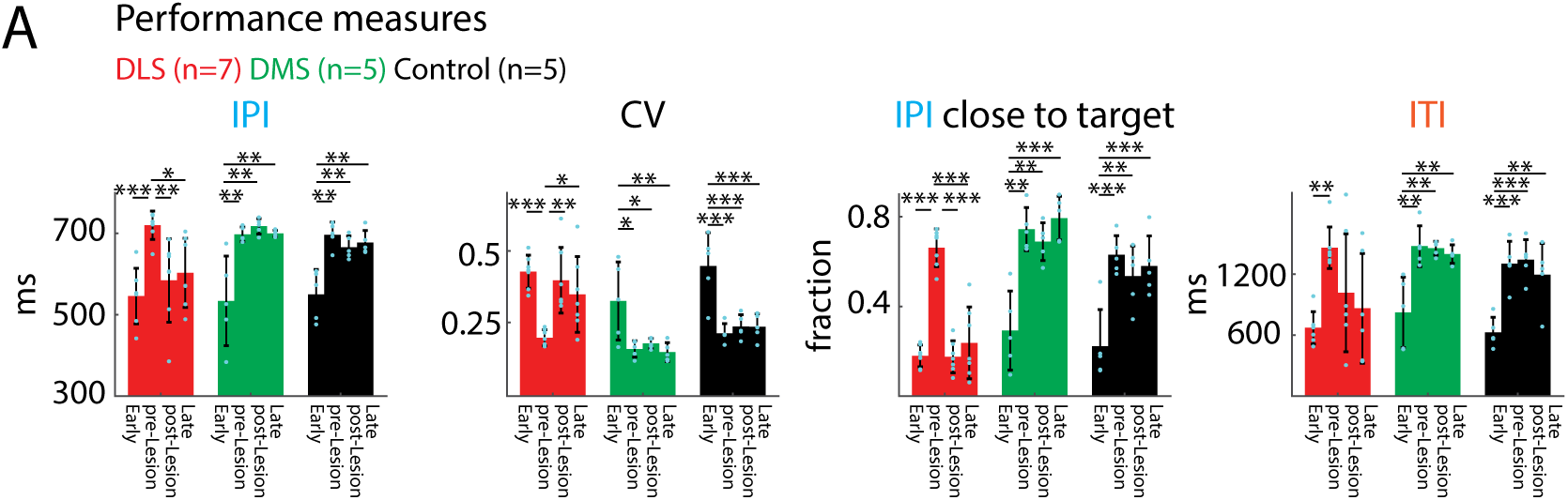
Task performance after DLS, but not DMS lesions is impaired, resembles performance early in training, and does not recover. **A.** Comparison of performance measures at different stages before and after manipulations (DLS n=7 rats; DMS n=5; Control n=5), showing long-lasting effects of DLS lesions and a reduction of performance to levels observed early in training. IPI: Inter Press Interval, CV of IPI: Coefficient of Variation of the IPI, IPI close to target: Fraction of trials close to target IPI (700 ms +/−20%), ITI: Inter Trial Interval. Early: First 2000 trials in training, pre-Lesion: First 2000 trials before lesion, post-Lesion: first 2000 trials after lesion, Late: trials 10,000 to 12,000 after lesion. Shown are means and error bars represent standard error of the mean (SEM). \**p* < 0.05, \*\**p* < 0.01, \*\*\**p* < 0.001.

**Supplementary Figure 5:**
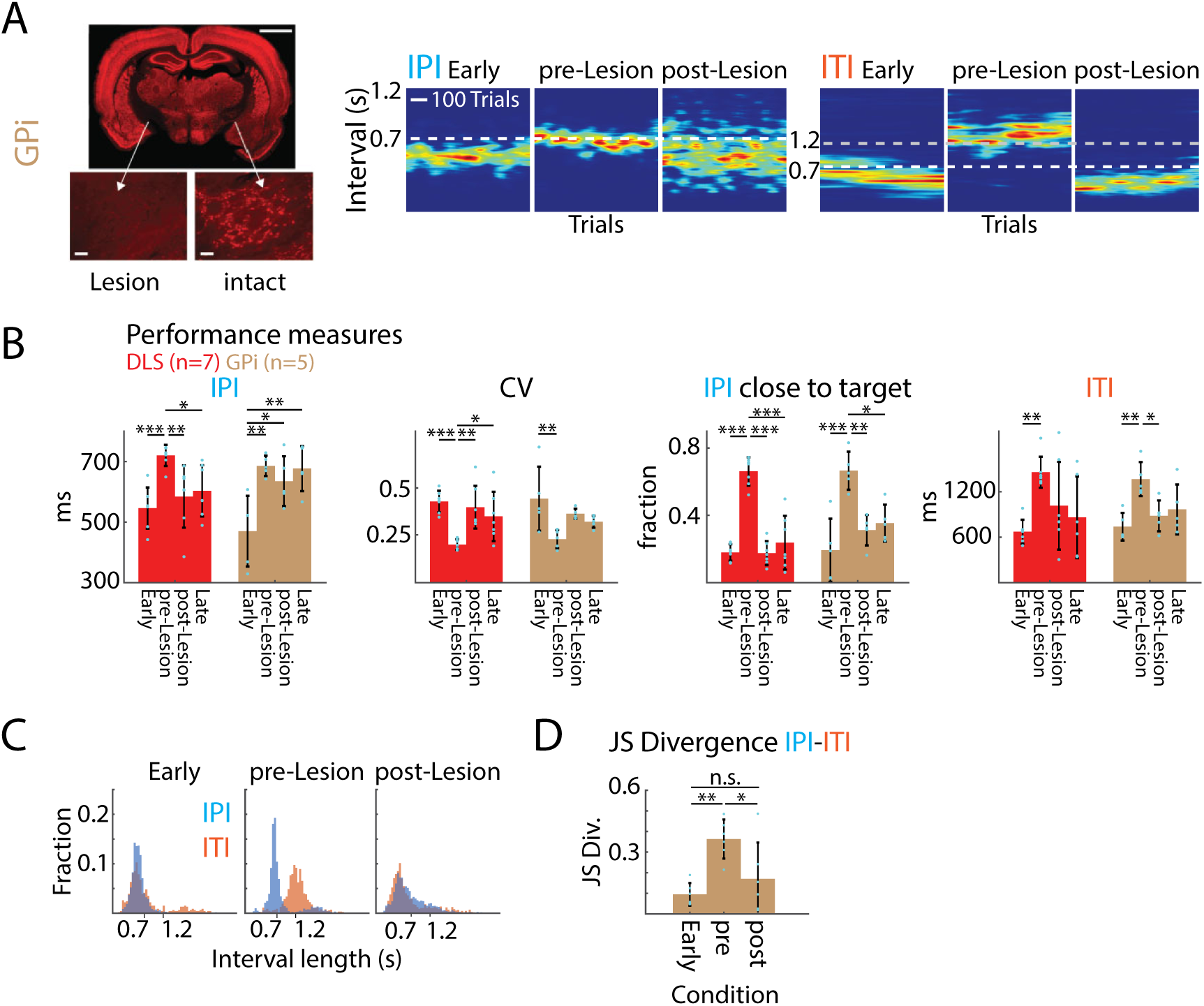
Lesions of the GPi / EP affect task performance similarly to DLS lesions. **A.** Representative example of the effect of a GPi lesion on task performance. Left: Example histological image of a unilateral GPi lesion, showing the comparison between the lesioned and the intact GPi. Experimental animals underwent bilateral GPi lesions (see Methods). Right: IPIs and ITIs for an example animal early in training, before and after bilateral GPi lesion. **B.** GPi lesions (n=5 rats) have long-lasting effects on various measures of performance (cf. S4 A). DLS performance as shown in Supplementary Figure 4, here shown for comparison. **C.** Left: Example distributions of IPI and ITI interval lengths early in training, and before and after GPi lesion. **D.** JS Divergence of IPI and ITI distributions for all GPi-lesioned animals (n=5 rats). Shown are means and error bars represent standard error of the mean (SEM). \**p* < 0.05, \*\**p* < 0.01.

## Methods

### Animals

The care and experimental manipulation of all animals were reviewed and approved by the Harvard Institutional Animal Care and Use Committee. Experimental subjects were female Long Evans rats 3-10 months old at the start of training (n=9 and n=28 for striatal recordings and manipulations, respectively; Charles River). Because the behavioral effects of our circuit manipulations could not be pre-specified before the experiments, we chose sample sizes that would allow for identification of outliers and for validation of experimental reproducibility. Animals were excluded from experiments post-hoc if the lesions were found to be outside of the intended target area or affected additional brain structures (see Lesion section). The investigators were not blinded to allocation during experiments and outcome assessment, unless otherwise stated.

### Behavioral Training

Rats were trained in a lever pressing task as previously described^31^. Water-restricted animals were rewarded with water for pressing a lever twice within performance-dependent boundaries around a prescribed interval between the presses (IPI = 700 ms). In addition, animals had to withhold pressing for 1.2 s after unsuccessful trials before initiating a new trial (inter-trial interval: ITI). All animals were trained in a fully automated home-cage training system^86^. Animals were only used for manipulations or recordings after they had reached our learning criteria (mean IPI = 700 ms +/−10%; CV of IPI distribution < 0.25 for a 3000-trial sliding window) and a median ITI > 1.2 s, indicating that they had learned the task structure and stabilized their performance.

### Surgical implantation and electrophysiological recordings

Surgical and recording procedures were as previously described^54^. Once rats reached asymptotic (expert) performance on the timed lever-pressing task, we performed surgery to implant microdrives containing arrays of 16 tetrodes into the dorsolateral striatum (n=3 rats) or dorsomedial striatum (n=3). In an additional cohort of animals (n=3), we performed recordings in the dorsolateral striatum after motor cortex lesion. For this, we performed two-stage bilateral lesions of motor cortex as previously described^31^ (see lesion surgeries below). During the surgery for the second motor cortex lesion, we also implanted the microdrive in the dorsolateral striatum. Animals were anesthetized with 1–3% isoflurane and placed in a stereotax. The skin was removed to expose the skull and five bone screws (MD-1310, BASi), including one soldered to a 200 µm diameter silver ground wire (786500, A-M Systems), were driven into the skull to anchor the implant. A stainless-steel reference wire (50 µm diameter, 790700, AM-Systems) was implanted in the external capsule to a depth of 2.5 mm, at a location posterior and contralateral to the implant site of the electrode array. After making a 4 to 5 mm diameter craniotomy and removing the dura, we slowly lowered the 16-tetrode array to a depth of 4.5 mm. Electrodes were targeted to a location 0.5 mm anterior and 4 mm lateral to bregma for dorsolateral striatum and 0.3 mm anterior and 2 mm lateral to bregma for dorsomedial striatum, and were implanted in the striatum contralateral to the dominant forelimb in the lever-pressing task. The microdrive was encased in a protective shield and cemented to the skull by applying a base layer of Metabond luting cement (Parkell) followed by a layer of dental acrylic (A-M systems).

After 7 days of recovery, rats were returned to their home-cages, which had been outfitted with an electrophysiology recording extension. The cage was placed in an acoustic isolation box, and training on the task resumed. Neural and behavioral data was recorded continuously for 12–16 weeks. Neural data was recorded using a 64-channel custom-designed head-stage (made of 2 RHD2132 ICs from Intan Technologies). The head-stage filtered, amplified, and digitized extracellular neural signals at 16 bits/sample and 30 kHz per second per channel. An FPGA module (Opal Kelly XEM6010 with Xilinx Spartan-6 FPGA) interfaced the head-stage with a computer that stored and displayed the acquired data. Behavioral data was acquired using high-speed imaging (at 120 Hz) from 2 cameras (Flea 3, Point Grey) placed on either side of the training cage. Video was synchronized to electrophysiological signals by recording TTL pulses from the CCD cameras that signaled frame capture times.

At the end of the experiments, animals were anesthetized and anodal current (30 µA for 30 s) passed through select electrodes to create micro-lesions at the electrode tips. Animals were transcardially perfused with phosphate-buffered saline (PBS) and subsequently fixed with 4% paraformaldehyde (PFA, Electron Microscopy Sciences) in PBS. Brains were removed and post-fixed in 4% PFA. Coronal sections (60 µm) were cut on a Vibratome (Leica), mounted, and stained with cresyl violet to reconstruct the location of implanted electrodes.

### Lesion surgeries

Bilateral striatal lesions, targeting either the motor cortex-recipient part (DLS) or the non-motor cortex input receiving part (DMS), and GPi (EP) lesions were performed in two stages. Once animals had reached asymptotic task performance (see Behavioral Training), the first striatal lesion was performed contralateral to the paw used for the first lever press in the acquired stereotyped motor sequence. After lesion and recovery (10 days), animals returned to training until their performance stabilized (at least 14 days after lesion). Subsequently, the ipsilateral striatal lesion was performed and after recovery animals were returned to training.

Lesions were performed as previously described^31^. Animals were anesthetized with 2% isoflurane in carbogen and placed in a stereotactic frame. After incision of the skin along the midline and cleaning of the skull, Bregma was located and small craniotomies for injections were performed above the targeted brain areas. A thin glass pipette connected to a micro-injector (Nanoject II, Drummond) was lowered to the injection site and an excitotoxin was injected. For striatal lesions quinolinic acid (0.09M in PBS (pH=7.3), Sigma-Aldrich) was injected in 4.9 nl increments to a total volume of 175 nl per injection site, at a speed of < 0.1 ul/min. For GPi lesions ibotenic acid (1% in 0.1M NaOH, Abcam) was injected in 4.9 nl increments to a total volume of 400 nl per injection site. After injection, the glass pipette was retracted by 100 um and remained there for at least 3 min before further retraction to allow for diffusion and to prevent backflow of the drug. After all injections were performed, the skin was sutured and animals received painkillers (Buprenorphine, Patterson Veterinary). Animals were allowed to recover for 10 days before being put back into training.

Injection coordinates were (in mm, according to Paxinos ^87^):

|  | DLS |  |  |  | DMS |  |  |  | GPi |
| --- | --- | --- | --- | --- | --- | --- | --- | --- | --- |
| AP* | +0.7 | +0.7 | -0.3 | -0.3 | +1.2 | +1.2 | +0.2 | +0.2 | -2.3 |
| ML* | +/-3.6 | +/-3.6 | +/-4.0 | +/-4.0 | +/-1.9 | +/-1.9 | +/-1.9 | +/-1.9 | +/-2.8 |
| DV | -5.5 | -4.5 | -5 | -4 | -5.5 | -4.5 | -5.5 | -4.5 | -7.5 |

\* measured from bregma

Motor cortex lesions for the cohort of animals in which we performed electrophysiological recordings in the dorsolateral striatum were performed analogous to striatal lesions and as previously described^31^. We injected ibotenic acid (1% in 0.1M NaOH, Abcam) in 4.9 nl increments to a total volume of 92 nl per injection site.

Injection coordinates were (in mm, according to Paxinos ^87^):

|  |  |  |  |  |  |  |
| --- | --- | --- | --- | --- | --- | --- |
|  | MC |  |  |  |  |  |
| AP* | +1 | +1 | +1 | +1 | +3 | +3 |
| ML* | +/-2 | +/-2 | +/-4.0 | +/-4.0 | +/-2 | +/-2 |
| DV | -1.6 | -0.8 | -1.6 | -0.8 | -1.6 | -0.8 |

\* measured from bregma

### Control surgeries

To test for nonspecific effects of surgery and striatal injections on behavior, we performed 1-stage control surgeries according to the procedure described above, bilaterally injecting different non-toxic solutions, either fluorophore-coated latex microspheres (red excitation [exc.] = 530 nm, emission [em.] = 590 nm and green exc. = 460 nm, em. = 505 nm) referred to as retrobeads (Lumafluor) ^88, 89^ or Adeno-associated viruses (AAVs) for non-specific expression of GFP (Penn Vector Core) into DLS. This allowed for post-hoc evaluation of the targeting of our control injections. Animals were returned to training after 10 days of recovery.

To determine which parts of the striatum receive input from either motor cortex (previously shown to be necessary for learning in our task^31^) or prefrontal Cortex (PFC) we injected AAVs for non-specific expression of GFP (Penn Vector Core) in either motor cortex or PFC. Injections were done in 9.2 nl increments, evenly spaced while slowly retracting the injection-pipette for a total volume of 300 nl per site / 1.5 ul per hemisphere for motor cortex and 500 nl per site / 1 ul per hemisphere for PFC. After surgery, we allowed for at least 4 weeks of viral expression before histological analysis (see Histology).

Injection coordinates were (in mm, according to Paxinos ^87^):

|  | MC |  |  |  |  | PFC |  |
| --- | --- | --- | --- | --- | --- | --- | --- |
| AP* | +1 | +1 | -1.5 | -2.25 | +3 | +3.2 | +2.2 |
| ML* | +/-2 | +/-4 | +/-2.75 | +/-2.5 | +/-2 | +/-0.6 | +/-0.6 |
| DV | -1.5 to -0.7 | -1.5 to -0.7 | -1.5 to -0.7 | -1.5 to -0.7 | -1.5 to -0.7 | -4.4 to -2.0 | -4.4 to -2.0 |

\* measured from bregma

### Histology

At the end of the experiment, animals were euthanized (100 mg/kg ketamine and 10 mg/kg xylazine), perfused with 4% paraformaldehyde (PFA), and their brains were harvested for histology to confirm lesion size and location or electrode implantation site. The brains were sectioned into 80 or 100-μm slices and stained with Cresyl Violet. In a subset of animals, immunofluorescence staining was performed instead of Cresyl Violet staining. After slicing, sections were blocked for 1h at room temperature in blocking solution (1% BSA, 0.3% TritonX), stained overnight at 4C with primary antibodies for NeuN (to stain for neuronal cell bodies; 1:500 in blocking solution; Millipore MAB377) and GFAP (to stain for glia cells; 1:500 in blocking solution; Sigma) and then with appropriate fluorescently-coupled secondary antibodies (1:1000 in blocking solution) for 2h at room temperature.

To determine the extent of the motor cortex and PFC projections in the striatum, immunofluorescent staining was performed in the same way, but instead of staining for GFAP, sections were stained for GFP to amplify the signal from the viral GFP expression.

Images of whole brain slices were acquired at 10x magnification with either a VS210 Whole Slide Scanner (Olympus) or an Axioscan Slide Scanner (Zeiss).

### Quantification of lesion size

To determine the extent and location of striatal lesions, we analyzed several sections (4-6) spanning the anterior-posterior extent of the striatum, allowing for an estimate of the overall lesion size. Lesion boundaries were determined throughout the striatum and adjacent areas, blind to the animals’ identity and performance. Boundaries were marked manually based on differences in cell morphology and density (loss of larger neuronal somata and accumulation of smaller glial cells). The extent of the striatum was determined based on the Paxinos Rat Brain Atlas^87^, using anatomical landmarks (external capsule, ventricle) and cell morphology and density. Additionally, we marked the GPe in posterior sections, since mistargeted injections may lead to its partial lesioning, disrupting the output both of the DLS and DMS. In addition to the overall lesion size, we also determined the lesioned fractions of the DLS/DMS. Since the DLS and DMS are not clearly defined, we made use of their differential input patterns from motor cortex and PFC, respectively, to estimate their extent. We used viral expression of GFP in motor cortex or PFC to visualize their respective axonal projection patterns in the striatum (n=3 each; see Control surgeries). Areas with axonal labeling in all animals were considered as motor cortex-input/PFC-input receiving. We then used these identified boundaries of the DLS and DMS to determine the lesioned fractions in the experimental animals. Based on these estimates we excluded animals with lesions affecting less than 50% of the respective target area or more than 10% of the non-targeted part of the striatum (n=4 rats). In addition, we excluded animals with lesions affecting a significant part of the GPe (>30%; n=3 rats).

### Kinematic Tracking

To determine the movement trajectories of the forelimbs and head of animals performing our task, we made use of recently developed machine learning approaches, using deep neuronal networks to determine the position of specific body parts in individual video frames^55^.

Videos of animals performing the task were acquired at 120 Hz by cameras pointing at the lever from either side and saved as snippets ranging from 1 s before the first lever press in a trial to 2 s after the last lever press in the trial. We extracted about 500 frames from each perspective randomly selected throughout the duration of the trials, balanced across pre- and post-manipulation conditions and manually labeled the position of the paws and head in each frame, using custom-written Matlab code. This data was used to train individual neural networks for each animal.

We trained ResNet-50 networks that were pretrained on ImageNet, using the original DeeperCut implementation in TensorFlow (https://github.com/eldar/pose-tensorflow)^55^. Training was performed using default parameters (1 million training iterations, 3 color channels, with pairwise terms, without intermediate supervision). Data augmentation was performed during training by rescaling images from a range of 85% to 115%.

The trained neural network was then used to predict the position of the body parts in all trials across conditions, on a frame-by-frame basis. The position of a body part in a frame is given by the peak of the network’s output score-map. Frames in which the body part was occluded were identified as having a low peak score. For both the training and the subsequent predictions we used GPUs in the Harvard Research Computing cluster.

Because the two forelimbs could often be confused for each other in the neural network’s predictions from a single frame, we took advantage of correlations across time to constrain the predictions. For each forelimb, the predicted score-maps for all frames in a single trial video were passed through a Kalman filter using the Python toolbox *filterpy*. Specifically, a constant-acceleration Kalman smoother was used which assumes that the forelimb on adjacent frames will have the same acceleration (zero jerk) plus a small noise term. Only frames with a weak neural-network prediction score were adjusted by the Kalman filter; otherwise the original neural-network prediction was used as the forelimb position.

The tracking accuracy was validated post-hoc by visual inspection of at least 50 predicted trajectories per animals. Initial training with lower frame numbers often led to inaccurate tracking results. After settling on a number of 500 training frames, none of the trained networks was discarded.

Missing frames in the trajectories, e.g. due to temporary occlusions of the forelimbs, were linearly interpolated for a maximum of 5 consecutive frames. If occlusions lasted longer, the trajectories were discarded. In a subset of animals, the quality of the recorded videos was not sufficient for high-quality tracking of the forelimbs, due to inappropriate lighting conditions or due to occlusions of the forelimbs over long durations of the trials and we had to discard the trajectories (n=1).

### Neural data analysis

We used our custom-designed spike-sorting algorithm fast automated spike tracker (FAST)^54^ to parse the raw neural data and isolate the spiking activity of populations of single units in an efficient and high-throughput manner. Units were separated into putative spiny projection neuron (SPN) and fast spiking interneuron (FSI) types using their spike-waveforms and average firing rates ^90^. Briefly, we performed k-means clustering (k=2) of units based on their peak-normalized spike-waveform, concatenated with their log-transformed firing rates.

For all analyses, we only considered neurons which had an average firing rate of at least 0.25 Hz within the period ranging from 1 s prior to and 2 s following the first lever press of the sequence. This eliminated 29%, 30%, and 42% of units recorded in the intact and motor cortex lesioned DLS, and the DMS, respectively.

### Peri-event time histograms (PETHs)

We computed peri-event time histograms (PETHs) of instantaneous firing rates aligned to the first lever press of the timed-lever pressing task. In order to restrict our analysis of neural activity to periods of stereotyped behavior, we selected only rewarded trials that followed previously rewarded trials (to control for the rat’s starting position), and these trials’ inter-press intervals had to be within 20% of the target inter-press interval of 700 ms (to ensure movement stereotypy). To account for the remaining variation in the length of the motor sequence, we linearly warped all spike-times that occurred between the 1^st^ and 2^nd^ lever press by a factor: *IPI*_t_⁄700 where *IPI*_t_ is the inter-press interval (in ms) on that trial^91^. When calculating PETHs, we computed the firing rate after binned the warped spike train within the period 1 s prior to and 2 s following the first lever press of the sequence, into 25 ms long bins. To calculate the Z-scored activity, we used a bootstrap approach to compute the mean and standard deviation of trial-averaged firing rates. We smoothed the Z-scored PETHs with a Gaussian kernel (σ = 25 ms) before plotting or calculating its maximum or minimum value, which we termed the Z modulation of that unit.

### Trial-by-trial correlations of neural activity

To calculate trial-by-trial correlations for a given neuron, we binned its spikes recorded within the period 1 s prior to and 2 s following the first lever press of the sequence into 25 ms bins on individual trials and then smoothed the spike-count vector with a Gaussian kernel (σ = 25 ms). We then calculated the correlation coefficients between the smoothed spike-count vectors for pairs of trials and averaged these measurements over all pairwise combinations of trials.

### Sparseness index

We calculated the sparseness index as previously described^92^. We calculated task-aligned histograms over trials of spikes recorded within the period 1 s prior to and 2 s following the first lever press of the sequence (25 ms bins). Histograms were normalized to calculate the spike probability within each bin (*p*_i_). We then computed the sparseness index (SI)^93^ as:

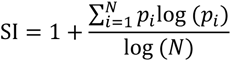

where *N* indicates the number of bins in the histogram. This index is 1 (maximal sparseness) when the activity is restricted to a single time bin and 0 if the spikes are evenly distributed across the time bins.

### Encoding analysis

We used generalized linear models (GLMs) to determine to what extent the instantaneous activity of striatal units could be predicted from the kinematics and timing of the motor sequence. For each unit, we calculated spike counts (25 ms bins) within the trial period ranging from 0.2 s before the 1^st^ lever press until 0.2 s after the 2^nd^ lever press. We excluded any units whose average firing rate within this period was less than 0.25 Hz. When fitting the GLMs, we used an exponential link function and modeled the observed spike counts with a Poisson distribution. 75% of the trials in each session were used for training the GLM, and the remaining 25% were held out for testing. We used elastic-net regularization (90% L1, 10% L2) to prevent over-fitting. The optimal value of the regularization penalty parameter (*λ*) was determined for each neuron separately using 5-fold cross-validation within the training set of trials. We used the software package “glmnet” for fitting the GLMs^94^.

We considered three sets of regressors – detailed kinematics of the forelimb and head movements, time within the sequence and vigor-related variables. (i) Kinematic features included the horizontal and vertical components of the velocity and acceleration of contra- and ipsilateral forelimbs as well as the head. These features were calculated after smoothing the raw trajectories using cubic spline interpolation (function *csaps* in Matlab, smoothing parameter = 0.1). The kinematic features were down-sampled by averaging to match the time-scale at which neural data was binned (25 ms bins). (ii) Time within the sequence was represented by a collection of vectors, each of which represented a Gaussian function (σ = 100 ms) whose mean was designated at a particular time within the template sequence (IPI = 700 ms). The means of the Gaussian functions for time vectors were uniformly spaced apart from their neighbors by 100 ms. This way the collection of time vectors constituted an overcomplete basis set to uniquely represent all times within the sequence for the GLM. These template vectors were then stretched or shrunk by interpolation to account for trial-by-trial deviations in the inter-press interval from the 700 ms template. (iii) To compute vigor-related variables, we measured the scalar magnitude of the vector velocity and acceleration of both forelimbs and the head, sampled at 25 ms intervals. We also used a combination of detailed kinematics and time within the sequence as an additional regressor set.

Goodness of fit of the encoding model was measured by a log-likelihood based measure termed the pseudo-R^2 95^.

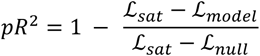

Where ℒ_model_ is the log-likelihood of observing spike-count data given the GLM’s predictions, ℒ_sat_ is the log-likelihood of a “saturated” model that has a parameter for each spike-count observation, and ℒ_null_ is the log-likelihood of the data given a “null” model that only fits the average spike count in the dataset.

### Decoding analysis

Following previous work^96^, we used a multilayer, feedforward neural network to predict the time-varying vertical and horizontal velocity components of the forelimbs and the head, or time within the sequence, using 75 ms of co-incident spiking activity (binned into 25 ms bins) of ensembles of striatal neurons. The network comprised two fully connected hidden layers of 400 units each with rectified linear activation functions. While training the network we used dropout on the 2 hidden layers (probability = 0.05). We used the Adam optimizer to train the neural network. We measured the accuracy of decoding using 4-fold cross-validation. For this analysis, we only considered behavioral sessions in which there were at least 10 simultaneously recorded units that had an average firing rate of at least 0.25 Hz within the trial period (from 0.2 s before and 0.2 s after the 1^st^ lever press and 2^nd^ lever press, respectively). For each of these sessions we fit decoding models using the activity of n=20 randomly sampled ensembles of size 4, 6, 8 and 10 striatal units. Decoding accuracy measured in each ensemble was then averaged across all 20 ensembles of the same size and then averaged across sessions for each rat.

### Behavioral Data Analysis

#### Performance Metrics

Performance metrics were determined based on the timing of lever presses in our task. The inter-press interval (IPI) was determined as the time between the first and second press in a trial, the inter-trial interval (ITI) as the time between the last press in an unsuccessful trial and the next occurring lever press. The CV was calculated across 25 trials and the moving average was low pass-filtered with a 50-trial boxcar filter. The fraction of trials close to the target IPI was calculated using the same windows and filters. Trials were labeled as close to the target if they were in the IPI range of 700 ms +/−20%.

#### JS Divergence

As a measure for the dissimilarity of the IPI and ITI distributions in individual animals, we calculated the Jensen-Shannon (JS) Divergence of the distributions. The JS Divergence is a symmetric derivative of the Kullback-Leibler divergence (KLD). We calculated the JS Divergence (JSD) as:

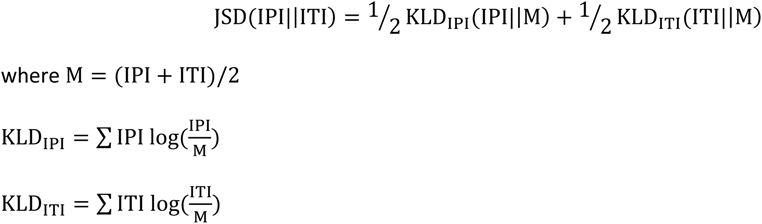

#### Trajectory analysis

We compared the trajectories of both forelimbs of all tracked animals before and after DLS or DMS lesions (Figures 6,7). We focused on the position of the forelimbs in the vertical dimension, in which the movements in our task are more pronounced than in the horizontal dimension. To be able to compare the stereotypy of the trajectories for the learned motor sequences, we sub-selected trials which were successful and rewarded, and which occurred after unrewarded trials. This allowed us to compare trials with the same start and end positions. This is necessary, since animals move down to, and back up from, a reward port underneath the lever after successful trials^31^. We plotted the average of the selected trajectories before and after the manipulation, calculated the SEM (Figure 6A,B) and plotted a projection of all selected trials (Figure 6A,B). To calculate the correlations between the individual trials, we linearly warped the trajectories to the same duration by interpolating between the lever presses. Since the lever presses themselves have stereotyped trajectories, largely independent of the trial duration, we interpolated only the trajectories from 100 ms after the first to 100 ms before the second lever press to preserve the shape of the presses. From these warped trajectories we calculated trial-to-trial correlations separately for both forelimbs and averaged the correlations for each trial (Figure 6A,B). These correlations were averaged for the individual conditions within animals and those means were averaged across animals and plotted with the SEM (Figure 6A,B).

To compare the trajectories across animals, we linearly warped all trajectories and normalized their amplitude to their individual maximum amplitude (Figure 6C,D). To calculate the correlations across animals, we first calculated the average pair-wise correlations across all trials within individual animals, and then averaged these across the individual animals (Figure 6C,D).

We separately compared the lever press movements, defined as the trajectory in the range of ±150 ms around a detected lever press (Figure 7A) and performed the same analysis as for the full trajectories in Figure 6. To compare the lever presses across animals before and after DLS lesion, we normalized the trajectories to their individual maximum amplitude and plotted their overlay (Figure 7B). As above, we calculated the average pairwise correlations for all lever presses in all trials of all animals across the conditions (pre- and post-lesion) and averaged them first by lever press (i.e. animal 1 press 1, animal 1 press 2, etc.) and then by condition (Figure 7B). To compare the lever presses after the lesion to the presses early in training, we additionally sub-selected trials as described above from the first 2000 trials of training. And performed the same analysis (Figure 7C).

To visualize the similarity between the individual lever presses, we performed a tSNE dimensionality reduction and plotted the coordinates of individual trials along the two resulting tSNE dimensions. For this we randomly sub-selected equal numbers of trials across conditions (Figure 7A, 2^nd^ column) and animals (Figure 7B,C, 2^nd^ column).

## Acknowledgements

We thank Sean Escola, James Murray and members of the Ölveczky lab for advice on data analysis and for discussions and comments on the manuscript. We thank Steve Turney and the Harvard Center for Biological Imaging for infrastructure and support. This work was supported by NIH grants R01-NS099323-01 and R01NS105349 to B.P.Ö., by a Life Sciences Research Foundation and Charles A. King Foundation postdoctoral fellowship to A.K.D. and by EMBO and HFSP postdoctoral fellowships to S.B.E.W.

## Author Contributions

A.K.D., S.B.E.W., R.K. and B.P.Ö. designed the study. A.K.D. and S.B.E.W. contributed equally and authorship order was determined by coin toss. A.K.D. performed electrophysiological recordings and analyzed the electrophysiology data. S.B.E.W. performed electrophysiological recordings, helped with the analysis and performed lesion experiments and analyzed behavioral data. R.K. performed pilot lesion experiments and described the initial phenomenology. A.K.D., S.B.E.W. and B.P.Ö. wrote the manuscript.

## References

1. Krakauer, J. W.. Hadjiosif, A. M.. Xu, J.. Wong, A. L..& Haith, A. M..Motor Learning. Compr. Physiol. 9, 613–663 (2019).

2. Grillner, S..& Robertson, B. The Basal Ganglia Over 500 Million Years. Curr. Biol. CB 26, R1088–R1100 (2016).

3. Grillner, S.. Robertson, B..& Stephenson-Jones, M. The evolutionary origin of the vertebrate basal ganglia and its role in action selection. J. Physiol. 591, 5425–5431 (2013).

4. Stephenson-Jones, M.. Samuelsson, E.. Ericsson, J.. Robertson, B..& Grillner, S. Evolutionary conservation of the basal ganglia as a common vertebrate mechanism for action selection. Curr. Biol. CB 21, 1081–1091 (2011).

5. Agostino, R.. Berardelli, A.. Formica, A.. Accornero, N..& Manfredi, M. Sequential arm movements in patients with Parkinson’s disease, Huntington’s disease and dystonia. Brain J. Neurol. 115 (**Pt 5**), 1481–1495 (1992).

6. Benecke, R.. Rothwell, J. C.. Dick, J. P.. Day, B. L..& Marsden, C. D. Disturbance of sequential movements in patients with Parkinson’s disease. Brain J. Neurol. 110 (**Pt 2**), 361–379 (1987).

7. Dudman, J. T..& Krakauer, J. W. The basal ganglia: from motor commands to the control of vigor. Curr. Opin. Neurobiol. 37, 158–166 (2016).

8. Graybiel, A. M. The Basal Ganglia and Chunking of Action Repertoires. Neurobiol. Learn. Mem. 70, 119–136 (1998).

9. Jin, X..& Costa, R. M. Shaping Action Sequences in Basal Ganglia Circuits. Curr. Opin. Neurobiol. 33, 188–196 (2015).

10. Turner, R. S..& Desmurget, M. Basal ganglia contributions to motor control: a vigorous tutor. Curr. Opin. Neurobiol. 20, 704–716 (2010).

11. Yttri, E. A..& Dudman, J. T. A Proposed Circuit Computation in Basal Ganglia: History-Dependent Gain. Mov. Disord. 33, 704–716 (2018).

12. Panigrahi, B..et al. Dopamine Is Required for the Neural Representation and Control of Movement Vigor. Cell 162, 1418–1430 (2015).

13. Rueda-Orozco, P. E..& Robbe, D. The striatum multiplexes contextual and kinematic information to constrain motor habits execution. Nat. Neurosci. 18, 453–460 (2015).

14. Turner, R. S.. Desmurget, M.. Grethe, J.. Crutcher, M. D..& Grafton, S. T. Motor Subcircuits Mediating the Control of Movement Extent and Speed. J. Neurophysiol. 90, 3958–3966 (2003).

15. Yttri, E. A..& Dudman, J. T. Opponent and bidirectional control of movement velocity in the basal ganglia. Nature 533, 402–406 (2016).

16. Anderson, M. E..& Horak, F. B. Influence of the globus pallidus on arm movements in monkeys. III. Timing of movement-related information. J. Neurophysiol. 54, 433–448 (1985).

17. Desmurget, M..& Turner, R. S. Testing Basal Ganglia Motor Functions Through Reversible Inactivations in the Posterior Internal Globus Pallidus. J. Neurophysiol. 99, 1057–1076 (2008).

18. Desmurget, M..& Turner, R. S. Motor Sequences and the Basal Ganglia: Kinematics, Not Habits. J. Neurosci. 30, 7685–7690 (2010).

19. Graybiel, A. M..& Grafton, S. T. The Striatum: Where Skills and Habits Meet. Cold Spring Harb. Perspect. Biol. 7, a021691 (2015).

20. Hikosaka, O. Role of Basal Ganglia in Control of Innate Movements, Learned Behavior and Cognition—A Hypothesis. in The Basal Ganglia IV: New Ideas and Data on Structure and function (eds. Percheron, G.. McKenzie, J. S. & Féger, J..) 589–596 (Springer US, 1994). doi:10.1007/978-1-4613-0485-2_61.

21. Mink, J. W. THE BASAL GANGLIA: FOCUSED SELECTION AND INHIBITION OF COMPETING MOTOR PROGRAMS. Prog. Neurobiol. 50, 381–425 (1996).

22. Barnes, T. D.. Kubota, Y.. Hu, D.. Jin, D. Z..& Graybiel, A. M. Activity of striatal neurons reflects dynamic encoding and recoding of procedural memories. Nature 437, 1158–1161 (2005).

23. Jin, X..& Costa, R. M. Start/stop signals emerge in nigrostriatal circuits during sequence learning. Nature 466, 457–462 (2010).

24. Martiros, N.. Burgess, A. A..& Graybiel, A. M. Inversely Active Striatal Projection Neurons and Interneurons Selectively Delimit Useful Behavioral Sequences. Curr. Biol. CB 28, 560–573.e5 (2018).

25. Thorn, C. A.. Atallah, H.. Howe, M..& Graybiel, A. M. Differential Dynamics of Activity Changes in Dorsolateral and Dorsomedial Striatal Loops during Learning. Neuron 66, 781–795 (2010).

26. Desrochers, T. M.. Amemori, K..& Graybiel, A. M. Habit Learning by Naive Macaques Is Marked by Response Sharpening of Striatal Neurons Representing the Cost and Outcome of Acquired Action Sequences. Neuron 87, 853–868 (2015).

27. Jin, X.. Tecuapetla, F..& Costa, R. M. Basal ganglia subcircuits distinctively encode the parsing and concatenation of action sequences. Nat. Neurosci. 17, 423–430 (2014).

28. Doyon, J.. Penhune, V..& Ungerleider, L. G. Distinct contribution of the cortico-striatal and cortico-cerebellar systems to motor skill learning. Neuropsychologia 41, 252–262 (2003).

29. Hikosaka, O..et al. Parallel neural networks for learning sequential procedures. Trends Neurosci. 22, 464–471 (1999).

30. Hikosaka, O.. Nakamura, K.. Sakai, K..& Nakahara, H. Central mechanisms of motor skill learning. Curr. Opin. Neurobiol. 12, 217–222 (2002).

31. Kawai, R..et al. Motor cortex is required for learning but not for executing a motor skill. Neuron 86, 800–812 (2015).

32. Grillner, S..& Robertson, B. The basal ganglia downstream control of brainstem motor centres — an evolutionarily conserved strategy. Curr. Opin. Neurobiol. 33, 47–52 (2015).

33. Lanciego, J. L.. Luquin, N..& Obeso, J. A..Functional neuroanatomy of the basal ganglia. Cold Spring Harb. Perspect. Med. 2, a009621 (2012).

34. McHaffie, J. G.. Stanford, T. R.. Stein, B. E.. Coizet, V..& Redgrave, P. Subcortical loops through the basal ganglia. Trends Neurosci. 28, 401–407 (2005).

35. Redgrave, P..& Coizet, V. Brainstem interactions with the basal ganglia. Parkinsonism Relat. Disord. 13 **Suppl 3**, S301–305 (2007).

36. Arber, S..& Costa, R. M. Connecting neuronal circuits for movement. Science 360, 1403–1404 (2018).

37. Grillner, S.. Hellgren, J.. Ménard, A.. Saitoh, K..& Wikström, M. A. Mechanisms for selection of basic motor programs – roles for the striatum and pallidum. Trends Neurosci. 28, 364–370 (2005).

38. Ruder, L..& Arber, S. Brainstem Circuits Controlling Action Diversification. Annu. Rev. Neurosci. 42, 485–504 (2019).

39. Aldridge, J. W..& Berridge, K. C. Coding of Serial Order by Neostriatal Neurons: A “Natural Action” Approach to Movement Sequence. J. Neurosci. 18, 2777–2787 (1998).

40. Aldridge, J. W.. Berridge, K. C..& Rosen, A. R. Basal ganglia neural mechanisms of natural movement sequences. Can. J. Physiol. Pharmacol. 82, 732–739 (2004).

41. Berridge, K. C..& Fentress, J. C. Deafferentation does not disrupt natural rules of action syntax. Behav. Brain Res. 23, 69–76 (1987).

42. Berridge, K. C..& Whishaw, I. Q. Cortex, striatum and cerebellum: control of serial order in a grooming sequence. Exp. Brain Res. Exp. Hirnforsch. Expérimentation Cérébrale 90, 275–290 (1992).

43. Cromwell, H. C..& Berridge, K. C. Implementation of action sequences by a neostriatal site: a lesion mapping study of grooming syntax. J. Neurosci. Off. J. Soc. Neurosci. 16, 3444–3458 (1996).

44. Markowitz, J. E..et al. The Striatum Organizes 3D Behavior via Moment-to-Moment Action Selection. Cell 174, 44–58.e17 (2018).

45. Roseberry, T. K..et al. Cell-Type-Specific Control of Brainstem Locomotor Circuits by Basal Ganglia. Cell 164, 526–537 (2016).

46. Takakusaki, K.. Saitoh, K.. Harada, H..& Kashiwayanagi, M. Role of basal ganglia–brainstem pathways in the control of motor behaviors. Neurosci. Res. 50, 137–151 (2004).

47. Gerfen, C. R..& Surmeier, D. J. Modulation of Striatal Projection Systems by Dopamine. Annu. Rev. Neurosci. 34, 441–466 (2011).

48. Graybiel, A. M. The basal ganglia: learning new tricks and loving it. Curr. Opin. Neurobiol. 15, 638–644 (2005).

49. Houk, J. C.. Adams, J. L..& Barto, A. G..A model of how the basal ganglia generate and use neural signals that predict reinforcement. in Models of information processing in the basal ganglia 249–270 (The MIT Press, 1995).

50. Hunnicutt, B. J..et al. A comprehensive excitatory input map of the striatum reveals novel functional organization. eLife 5, e19103 (2016).

51. Mandelbaum, G..et al. Distinct Cortical-Thalamic-Striatal Circuits through the Parafascicular Nucleus. Neuron 102, 636–652.e7 (2019).

52. Alexander, G. E.. DeLong, M. R..& Strick, P. L. Parallel Organization of Functionally Segregated Circuits Linking Basal Ganglia and Cortex. Annu. Rev. Neurosci. 9, 357–381 (1986).

53. Hintiryan, H..et al. The mouse cortico-striatal projectome. Nat. Neurosci. 19, 1100–1114 (2016).

54. Dhawale, A. K..et al. Automated long-term recording and analysis of neural activity in behaving animals. eLife 6, e27702 (2017).

55. Insafutdinov, E.. Pishchulin, L.. Andres, B.. Andriluka, M..& Schiele, B..DeeperCut: A Deeper, Stronger, and Faster Multi-person Pose Estimation Model. in Computer Vision – ECCV 2016 (eds. Leibe, B.. Matas, J.. Sebe, N. & Welling, M.) 34–50 (Springer International Publishing, 2016).

56. Mathis, A..et al. DeepLabCut: markerless pose estimation of user-defined body parts with deep learning. Nat. Neurosci. 21, 1281–1289 (2018).

57. Yin, H. H..et al. Dynamic reorganization of striatal circuits during the acquisition and consolidation of a skill. Nat. Neurosci. 12, 333–341 (2009).

58. Vandaele, Y..et al. Distinct recruitment of dorsomedial and dorsolateral striatum erodes with extended training. eLife 8, e49536 (2019).

59. Sales-Carbonell, C..et al. No Discrete Start/Stop Signals in the Dorsal Striatum of Mice Performing a Learned Action. Curr. Biol. 28, 3044–3055.e5 (2018).

60. Hilario, M.. Holloway, T.. Jin, X..& Costa, R. M. Different dorsal striatum circuits mediate action discrimination and action generalization. Eur. J. Neurosci. 35, 1105–1114 (2012).

61. Yin, H. H. The Sensorimotor Striatum Is Necessary for Serial Order Learning. J. Neurosci. 30, 14719– 14723 (2010).

62. Hammond, C.. Bergman, H..& Brown, P. Pathological synchronization in Parkinson’s disease: networks, models and treatments. Trends Neurosci. 30, 357–364 (2007).

63. Jenkinson, N..& Brown, P. New insights into the relationship between dopamine, beta oscillations and motor function. Trends Neurosci. 34, 611–618 (2011).

64. Marreiros, A. C.. Cagnan, H.. Moran, R. J.. Friston, K. J..& Brown, P. Basal ganglia-cortical interactions in Parkinsonian patients. NeuroImage 66, 301–310 (2013).

65. Uhlhaas, P. J..& Singer, W. Neural Synchrony in Brain Disorders: Relevance for Cognitive Dysfunctions and Pathophysiology. Neuron 52, 155–168 (2006).

66. Lozano, C. S.. Tam, J..& Lozano, A. M. The changing landscape of surgery for Parkinson’s Disease. Mov. Disord. 33, 36–47 (2018).

67. Okun, M. S..& Vitek, J. L. Lesion therapy for Parkinson’s disease and other movement disorders: Update and controversies. Mov. Disord. 19, 375–389 (2004).

68. Dudman, J. T..& Krakauer, J. W. The basal ganglia: from motor commands to the control of vigor. Curr. Opin. Neurobiol. doi:10.1016/j.conb.2016.02.005.

69. Redgrave, P.. Prescott, T. J..& Gurney, K. The basal ganglia: a vertebrate solution to the selection problem? Neuroscience 89, 1009–1023 (1999).

70. Esposito, M. S.. Capelli, P..& Arber, S. Brainstem nucleus MdV mediates skilled forelimb motor tasks. Nature 508, 351–356 (2014).

71. Humphries, M. D.. Gurney, K..& Prescott, T. J. Is there a brainstem substrate for action selection? Philos. Trans. R. Soc. Lond. B. Biol. Sci. 362, 1627–1639 (2007).

72. Geddes, C. E.. Li, H..& Jin, X. Optogenetic Editing Reveals the Hierarchical Organization of Learned Action Sequences. Cell 174, 32–43.e15 (2018).

73. Aldridge, J. W.. Berridge, K. C.. Herman, M..& Zimmer, L. Neuronal Coding of Serial Order - Syntax of Grooming in the Neostriatum. Psychol. Sci. 4, 391–395 (1993).

74. Grillner, S..& Wallén, P. Innate versus learned movements--a false dichotomy? Prog. Brain Res. 143, 3–12 (2004).

75. Haber, S. N..Corticostriatal circuitry. Dialogues Clin. Neurosci. 18, 7–21 (2016).

76. McGeorge, A. J..& Faull, R. L. The organization of the projection from the cerebral cortex to the striatum in the rat. Neuroscience 29, 503–537 (1989).

77. Murray, J. M..& Escola, G. S. Learning multiple variable-speed sequences in striatum via cortical tutoring. eLife 6, e26084 (2017).

78. Hazeltine, E.. Helmuth, L. L..& Ivry, R. B. Neural mechanisms of timing. Trends Cogn. Sci. 1, 163–169 (1997).

79. Ivry, R. B. The representation of temporal information in perception and motor control. Curr. Opin. Neurobiol. 6, 851–857 (1996).

80. Freeman, J. S..et al. Abnormalities of motor timing in Huntington’s disease. Parkinsonism Relat. Disord. 2, 81–93 (1996).

81. Harrington, D. L.. Haaland, K. Y..& Hermanowicz, N. Temporal processing in the basal ganglia. Neuropsychology 12, 3–12 (1998).

82. Fee, M. S..& Scharff, C. The songbird as a model for the generation and learning of complex sequential behaviors. ILAR J. 51, 362–377 (2010).

83. Hahnloser, R. H. R.. Kozhevnikov, A. A..& Fee, M. S. An ultra-sparse code underlies the generation of neural sequences in a songbird. Nature 419, 65–70 (2002).

84. Simpson, H. B..& Vicario, D. S. Brain pathways for learned and unlearned vocalizations differ in zebra finches. J. Neurosci. Off. J. Soc. Neurosci. 10, 1541–1556 (1990).

85. Long, M. A..& Fee, M. S. Using temperature to analyse temporal dynamics in the songbird motor pathway. Nature 456, 189–194 (2008).

86. Poddar, R.. Kawai, R..& Ölveczky, B. P. A Fully Automated High-Throughput Training System for Rodents. PLoS ONE 8, e83171 (2013).

87. Paxinos, G..& Watson, C. A stereotaxic atlas of the rat brain. N. Y. Acad. (1998).

88. Katz, L. C..& Iarovici, D. M. Green fluorescent latex microspheres: a new retrograde tracer. Neuroscience 34, 511–520 (1990).

89. Katz, L. C.. Burkhalter, A..& Dreyer, W. J. Fluorescent latex microspheres as a retrograde neuronal marker for in vivo and in vitro studies of visual cortex. Nature 310, 498–500 (1984).

90. Berke, J. D.. Okatan, M.. Skurski, J..& Eichenbaum, H. B. Oscillatory Entrainment of Striatal Neurons in Freely Moving Rats. Neuron 43, 883–896 (2004).

91. Leonardo, A..& Fee, M. S. Ensemble Coding of Vocal Control in Birdsong. J. Neurosci. 25, 652–661 (2005).

92. Ölveczky, B. P.. Otchy, T. M.. Goldberg, J. H.. Aronov, D..& Fee, M. S. Changes in the neural control of a complex motor sequence during learning. J. Neurophysiol. 106, 386–397 (2011).

93. Lehky, S. R.. Sejnowski, T. J..& Desimone, R. Selectivity and sparseness in the responses of striate complex cells. Vision Res. 45, 57–73 (2005).

94. Friedman, J. H.. Hastie, T..& Tibshirani, R. Regularization Paths for Generalized Linear Models via Coordinate Descent. J. Stat. Softw. 33, 1–22 (2010).

95. Colin Cameron, A. & Windmeijer, F. A. G. An R-squared measure of goodness of fit for some common nonlinear regression models. J. Econom. 77, 329–342 (1997).

96. Glaser, J. I..et al. Machine learning for neural decoding. ArXiv170800909 Cs Q-Bio Stat (2019).

